# Gradient in cytoplasmic pressure in the germline cells controls overlying epithelial cell morphogenesis

**DOI:** 10.1101/440438

**Authors:** Laurie-Anne Lamiré, Pascale Milani, Gaël Runel, Annamaria Kiss, Leticia Arias, Blandine Vergier, Pradeep Das, David Cluet, Arezki Boudaoud, Muriel Grammont

## Abstract

It is unknown how growth in one tissue impacts morphogenesis in a neighboring tissue. To address this, we used the *Drosophila* ovarian follicle, where a cluster of 15 nurse cells and a posteriorly located oocyte are surrounded by a layer of epithelial cells. It is known that as the nurse cells grow, the overlying epithelial cells flatten in a wave that begins in the anterior. Here, we demonstrate that an anterior to posterior gradient of decreasing cytoplasmic pressure is present across the nurse cells and that this gradient acts through TGFß to control both the triggering and the progression of the wave of epithelial cell flattening. Our data indicate that intrinsic nurse cell growth is important to control proper nurse cell pressure. Finally, we reveal that nurse cell pressure and subsequent TGFß activity in the StC combine to increase follicle elongation in the anterior, which is crucial for allowing nurse cell growth and pressure control. More generally, our results reveal that during development, inner cytoplasmic pressure in individual cells has an important role in shaping their neighbors.

**Impact Statement:** Cell shape change depends on extrinsic forces exerted by cytoplasmic pressure in neighbouring cells.

## Introduction

Epithelial cells collectively adopt specific shapes when building organs. Over the last four decades, it has been demonstrated that cell and organ morphogenesis depend on the integration of extrinsic and intrinsic biological cues (endocrine, paracrine or autocrine signaling; cell-cell or cell-extracellular matrix adhesion; actin filament and microtubule organization etc.). Internal forces generated by the activity of the acto-myosin network has been shown to be crucial for cell shape, by acting on adherens junction and/or on cell-extracellular matrix adhesive complexes [1,2,11– 18,3–10]. Such studies shed light on the importance of considering the mechanical properties of cells, such as cortical tension, to understand morphogenesis [19–22]. However, cell shape is not only imposed by intrinsic forces, it also depends on the local environment, which comprises other types of cells and extra-cellular matrix. The mechanical properties of these biological elements may also impact and deform epithelial cells. The implication in epithelial morphogenesis of such extrinsic forces is far less investigated and known, compared to intrinsic forces [23, 24].

With its simple structure and its well-characterized pattern of development, the *Drosophila* ovarian follicle is a valuable model to study epithelial cell morphogenesis. A follicle consists of an inner cyst of 16 germinal cells (15 nurse cells and one posteriorly-localized oocyte) surrounded by a monolayered epithelium of about 800 cells, which are themselves covered by an outer basement membrane (BM) (Figure 1A). Follicle development has been divided in 14 stages with stages 1 and 14 corresponding to a follicle emerging from the germarium and a mature egg, respectively [25, 26]. The follicle is initially small and spherical with a 10 µm diameter. It progressively grows up and elongates before undergoing a major acceleration of growth and elongation at stage 9. In parallel, the oocyte is gradually enlarged compared to individual nurse cell (NC), due to the presence of ring canals (RC) between all the germline cells that serve to transfer NC cytoplasmic contents into the oocyte [27]. The epithelial cells accommodate the germline growth by first proliferating from S1 to S6 to reach about 850 cells and second by increasing in volume and changing in shape at stage 9 [25, 26]. Starting from a cuboidal shape, about fifty cells, called the stretched cells (StC), flatten above the nurse cells while the others, called the main body follicular cells (MBFC), become columnar above the oocyte. StC flattening occurs progressively from the anterior to the middle of the follicle and can be monitored by following the increase of the apical surface (Figure 1B). This morphogenetic process depends on the activation of specific genes in the StC, such as those involved in the TGFß pathway and the *tao* gene, which control the sterotyped disassembly of the sub-apical adherens junctions (AJ) (Figure 1B’) and the lateral adhesion complexes, respectively [28–30]. Cell flattening also comes with changes in BM mechanical properties and interactions, which are both controlled by the TGFß pathway [31]. Finally, StC flattening depends on the growth of the germline cells that controls the degree of flattening [32]. How this external force influences StC flattening has never been investigated.

**Figure 1:**
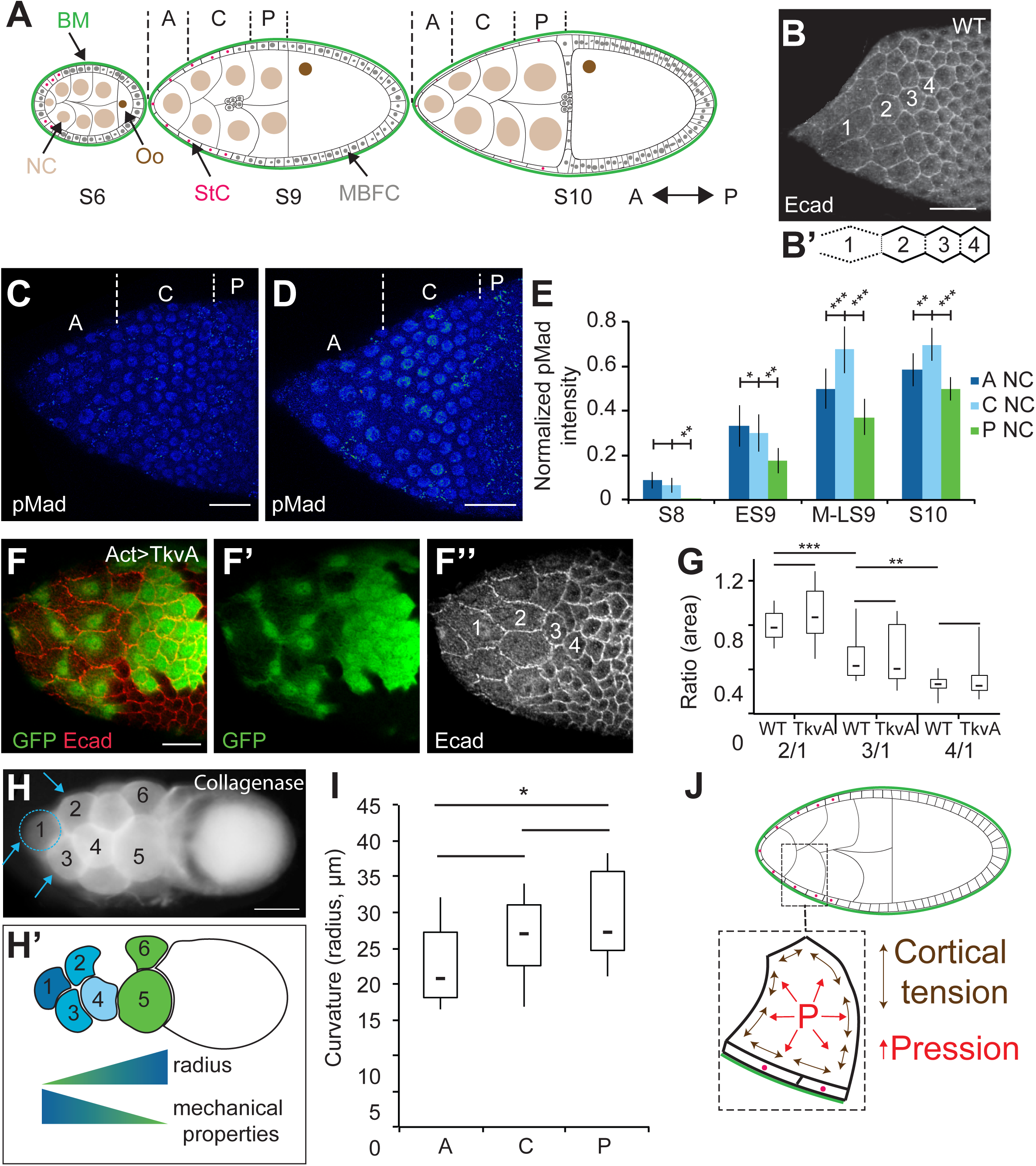
An A/P gradient of mechanical properties is present in the nurse cells. In all figures, anterior is to the left. (A) Schematic representation of three follicles at different developmental stages (S6, S9 and S10) with nurse cells (NC), oocyte (Oo), main body follicular cells (MBFC), stretched cells (StC) and basement membrane (BM) indicated. From S8 onwards, NC are grouped into three regions for analysis: anterior (A), central (C) and the four posterior (P) NC. (B) Adherens junctions (Ecad) remodelling and increase of apical surface occur in the StC of a S9 follicle. Both parameters are marker of flattening (see movie 1). Four cells are labelled from anterior (1) to posterior (4) to highlight changes in apical surface. (B’) Schematic representation of AJ remodelling, with dotted lines representing remodelled AJ and solid lines representing intact AJ. The AJ disassembly consists of first remodeling the vertices on the same row and the AJ perpendicular to the antero-posterior axis (A/P), second elongating the AJ parallel to the AP axis, and third, disassembling those AJ (Figure 1B’). (C, D) pMad expression in early (C) and mid (D) WT S9 follicles. The anterior (A), central (C) and posterior (P) regions of the NC are indicated. (E) Quantification of pMad in follicular cells during StC flattening (n > 15 nuclei per condition). (F) TkvA-expressing StC in a S9 follicle marked by GFP (F and F’), and with StC labelled in decreasing order from anterior to posterior (F’’). (G) Box and whisker plot of the ratio of surface area of StC in WT or TkvA-expressing StC at the different positions shown in B or F’’, respectively (n > 23 cells per plot). Scale bars: 20 µm. (H) A WT S10 follicle after collagenase treatment, showing the NC bulging outwards, with the corresponding schematic representation of the mechanical property gradient based on NC membrane curvature (the measured curvature corresponds to the membrane bulging outwards) (H’). (I) Box and whisker plot of the radius of curvature of anterior (A), central (C) or posterior (P) NC (n > 20 per region) in late S9 and S10 follicles. (J) Schematic representation of a stage 9 follicle, with a blow up on a nurse cell, its overlying StC and the BM. In the NC, cortical tension (double sided arrows, in brown) and cytoplasmic pressure (P, in red) are represented. Scale bars: 20 µm (B-F) and 50 µm (H). In box and whisker plots in all figures, boxes extend from 25 to 75 percentile, with a line showing the median value. Whiskers extend to the most extreme values. In all figures, error bars indicate s.e.m. In all figures, *, ** and *** correspond to p<0.5, p<0.05 and p<0.01 (t-test), respectively.

In this study, we establish that a gradient of mechanical properties exists in the NC compartment along the anterior-posterior axis and that this gradient is responsible for the progressive change of shape of the StC. We show that the differences in NC mechanical properties to differences in cytoplasmic pressure, which is the force that counters cortical tension. We present evidence that pressure level is modulated by intrinsic NC growth and controls TGFß expression dynamics. We reveal that NC pressure leads to inhomogeneous follicle expansion, with a bias toward the anterior and that this expansion requires StC flattening and BM softening. We show that anterior expansion is crucial for the maintenance of NC integrity while their inner pressure builds up. Whereas virtually no study has paid attention to cytoplasmic pressure as a physical component of the cell, our data bring to light its role in imposing shape in epithelial cells and tissues.

## Results

### The wave of StC flattening does not depend solely on TGFß activity

StC flattening occurs progressively from anterior to posterior and depends on TGFß signaling (Brigaud et al., 2015). To determine whether the wave of flattening depends on a wave of TGFß activity, we quantified the expression of the phosphorylated form of Mad (pMad) from stage 8 to stage 10. pMad is mainly detected in epithelial cells above the anterior and central NC with the highest levels in the anterior cells. Only a weak expression is detected in the epithelial cells surrounding the posterior NC. As StC flattening progresses posteriorly, pMad level increases in all the epithelial cells that surround NC, with a maximum detected in those above the central part. These data confirm that TGFß activity levels vary in the cells covering the NC and that these variations follow the progression of StC flattening (Figure 1C - 1E) [33]. To determine the importance of the spatio-temporal pattern of TGFß for the progressive change of shape, we measured StC apical surface area when expressing a constitutively active form of the TGFß receptor Thick veins (Tkv), which we refer to as TkvA. Our data show that the TkvA-expressing StC still flatten like a wave from anterior to posterior, demonstrating that cell flattening remains progressive in the absence of graded TGFß activity (Figure 1F, 1G). We conclude that another parameter, independent from the TGFß pathway, must therefore control the wave of flattening.

### An antero-posterior gradient of mechanical properties is present in the nurse cell compartment

We previously observed that a collagenase treatment on follicles leads to the bulging of the NC toward the outside, when the constraint imposed by the basement membrane is removed (Figure 1H) [31]. This also shows that StC are insufficient to constrain NC under collagenase treatment. Furthermore, this experimental condition facilitates the assessment NC mechanical properties. Cell mechanical properties are made up of mainly two components: cortical tension, controlled by the cortical acto-myosin network and by cell-cell adhesion, and the opposed cytoplasmic pressure, which may vary with solute and water fluxes, with differences in volume or growth rate, and with organization of cytoplasmic structural elements, such as microtubules and non-cortical actin microfilaments. We reasoned that the radius of the circle fitting the outward-facing NC membrane (the radius of curvature) may reflect its NC mechanical properties, given that the radius of curvature (R) of an interface is proportional to the ratio of its tension (γ) and inversely proportional to the difference in pressure (ΔP) between the two sides of the interface: R = 2γ / ΔP [34, 35]. We therefore measured NC radii along the A/P axis and found a gradient of curvature with smaller radii at the anterior in late S9 and S10 follicles (Figure 1H, 1I), indicating that the NC mechanical properties vary along the A/P axis, with tension increasing or pressure decreasing from anterior to posterior, or both (Figure 1J).

### The A/P gradient of NC mechanical properties is consistent with differences in cortical tension at late stage 9

To test whether the observed variations in NC curvature from anterior to posterior could be due to increasing NC surface tension, we considered surface tension of the NC during StC flattening in follicles with intact basement membrane. The tension (γ) of a cell-cell interface is increased by contractility of the actomyosin cortex (γ_am_) and reduced by cell-cell adhesion (γ_ad_), and can be stated as: γ = γ_am_ - γ_ad_ [36–38]. Accordingly, we analyzed cortical actomyosin dynamics through the expression and activity of the regulatory chain of the non-muscular Myosin II (encoded by *spaghetti squash*, *sqh*) and cell-cell adhesion through the expression of Ecadherin (Ecad), in live and fixed follicles from stage 8 to stage 10 (Figure 2A and Figure S1A – S1C). Ecad and Sqh (both the non-phosphorylated and the phosphorylated, pSqh, forms) are strongly detected at the cortex of the oocyte from stage 8 to 10. Ecad expression is weak in the NC at stage 8 and increases during stage 9. Sqh expression is undetectable at stage 8 but starts to be visible at stage 9. In stage 9 follicles, pSqh is also detected at a low level at the NC cortex. To quantify and compare Ecad, Sqh and pSqh expressions along the A/P axis, expression levels at the NC membranes along the A/P axis were normalized for each follicle with that of the oocyte cortex. The anterior NC usually express a higher Ecad level than the central NC from mid stage 9 to stage 10, indicating higher adhesion in the anterior, negatively influencing tension there. Sqh levels remain constant at all NC interfaces, regardless of position, throughout these stages, though the active form of Sqh does show an inverse gradient of expression with higher levels in central and posterior cells compared to anterior cells at the end of stage 9, indicating lower contractility at the anterior at this stage. Altogether, these observations suggest that a postero-anterior gradient of cortical tension is present from mid to late stage 9.

**Figure 2.**
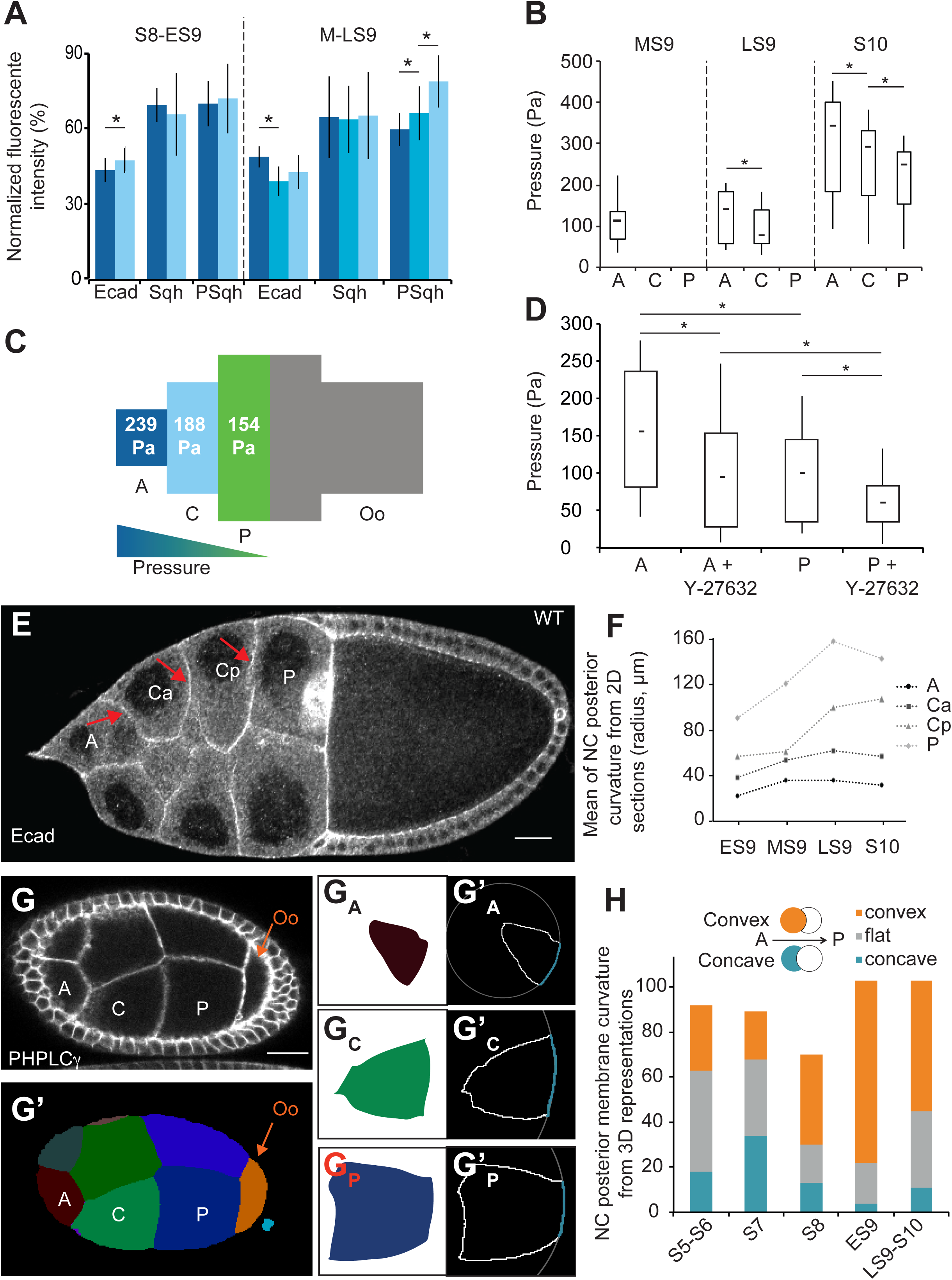
AFM measurements and nurse cell geometry reveal an A/P gradient of cytoplasmic pressure. (A) Percentage of Ecad::GFP (live follicles), Sqh::RFP (live follicles), and pSqh (fixed follicles) expression at the interface between anterior and central (dark blue) NC, or between central abuting the anterior NC and central abuting the posterior NC (Cyan, only at mid and late stage 9), or between central and posterior (light blue) NC, relative to the expression between posterior NC and the oocyte (n>15 per stage). (B) Box and whisker plot of cytoplasmic pressure in anterior (A), central (C) and posterior (P) NC, from mid S9 (MS9) to S10 (S10) WT follicles (n > 10 per bar per stage). (C) Schematic representation of mean cytoplasmic pressure in each region of WT S10 follicles, with a color-coded gradient from blue (high) to green (low). (D) Box and whisker plot of cytoplasmic pressure in anterior (A) and posterior (P) NC, at mid S9 WT follicles before or after treatment with ROCK inhibitor (Y-27632), (n = 13 per bar). (E) A late S9 follicle showing the curvature of the membranes (arrows) between the anterior (A) and a central NC (Ca), between two central NC (Ca and Cp), or between a central (Cp) and a posterior (P) NC. Ca and Cp refer to a central NC in contact with an anterior NC or a posterior NC, respectively. (F) Mean of the NC posterior membrane curvatures (as labelled in E) at either early-S9 (ES9; n = 98 cells), mid S9 (MS9; 42 cells), late S9 (LS9; 25 cells) or S10 (S10; 15 cells). (G) A S8 follicle and the corresponding section from a 3D segmentation of the same (G’) with NC labelled from anterior to posterior. G_A_, G_C_ and G_P_ correspond to slices showing the largest area for each labelled NC. G’_A_, G’_C_ and G’_P_ show the outline (solid white line) of each cell, as well as the circle (dotted white line) fitted to the posterior membrane (blue line). (H) Number of convex (orange), flat (grey) and concave (blue) NC posterior membrane curvatures at different follicular stages (2 - 3 follicles per stage were reconstructed). Scale bars: 20 µm

### The antero-posterior gradient of NC mechanical properties corresponds to differences in cytoplasmic pressure

To test whether the observed variations in NC curvature from anterior to posterior could be due to decreasing NC cytoplasmic pressure, we used an atomic force microscope (AFM), a standard approach to non-invasively measure the stiffness of biological samples [39–45]. The apparent stiffness quantifies how easily a body is deformed when a force is applied to it; a high value corresponds to a stiff material. We designed a bespoke AFM-based protocol with deep sample indentation so as to probe NC internal pressure below the basement membrane and the flattened StC (see Methods, Figure S1D - S1F). From the force curves, we first quantified the apparent stiffness along the follicle (Figure S1G). In parallel, we also quantified the curvature of the follicle surface at the measurement points. With the values of indentation depth used, we extracted pressure using stiffness and curvature values (Figure S1H) (see Methods), as previously described [45]. We observed that inner pressure of anterior NC increases by a factor of 3 during StC flattening and that an antero-posterior gradient of pressure is present, with the highest pressure being in the most anterior probed area (Figure 2B, 2C and Figure S2A). Although NC pressure displays a broad range of values, between 100 Pa and 450 Pa, we observe that it varies by a factor of 1.2- to 1.5-fold between different regions of a single follicle (Figure S2A), whereas variations within each probed region were negligible (Figure S2B, S2C). We note that the values of pressure measured are comparable to those obtained in mitotic cells [46], chick embryo blastula (Henkels et al., 2013) and in mouse blastocyst lumen [48].

Although we chose an indentation depth that is more sensitive to NC cytoplasmic pressure, it is possible that these measurements are also sensitive to NC cortical tension. In order to assess this, we performed measurements (n=13) before and after the addition of a pharmacological ROCK inhibitor (Y-27632), which has been shown to inhibit Myosin activity (He et al., 2010). We found that blocking Myosin II decreases pressure by a factor of 1.6 for both anterior and posterior NC (Figure 2D). Consequently, the ratio in stiffness between anterior and posterior NC is preserved after ROCK inhibitor treatment, demonstrating that these differences are mostly due to cytoplasmic pressure and not to cortical tension. In addition, we performed an osmotic treatment by adding 1M NaCl to the medium and observed a sharp decrease in stiffness values (n=8), demonstrating that measurements are responsive to osmotic changes (Figure S2D, S2E). Taken together, these data demonstrate the presence of a cytoplasmic pressure gradient within the NC compartment at late stage 9 and at stage 10.

Since curvature of a soft interface is physically ascribed to differences in hydrodynamic pressure between the two sides of this interface [34, 35], we predicted from the gradient in pressure that the interfaces between NC should be curved towards the posterior in normal conditions. To test this, we examined NC membrane shape and quantified the curvatures of NC membranes in fixed tissues throughout stage 9 (Figure 2E, 2F and Figure S3A, S3B). This analysis shows that NC membranes bulge toward the posterior while StC are flattening and that a gradient of curvature is observed from anterior to posterior NC (those abutting the oocyte), with the anterior NC being the most curved. We also analyzed membrane curvature in follicles treated with the ROCK inhibitor and found that most of the membranes still bulge toward the posterior, ruling out a putative role of cortical tension in this gradient of curvature (Figure S3C, S3D). Finally, to generate more precise data on membrane curvature, we generated 3D reconstructions of live follicles (Movie 1). To this end, we used the MARS pipeline (Multi-Angle image acquisition, three-dimensional Reconstruction and cell Segmentation) [49] (Figure S3E - S3H), and additionally developed an ImageJ macro to automatically measure membrane curvature in individual NC (see Methods, Figure S4A). We observed that posterior NC membranes become progressively convex from S7 to S9, and that this is correlated with anterior NC membranes becoming concave (Figure 2G, 2H and Figure S4B, S4C, S4E). At S9, 72 % of posterior-facing membranes are convex compare to 3% of anterior-facing membranes, while no specific orientation is ever observed for the lateral membranes (Figure S4F), consistent with the 2D measurements. Finally, we also noticed that the anterior membranes of the oocyte first bulge into the posterior NC (convex) up to stage 7 before switching to being concave during stage 8 (Figure S4D - S4G, Movie 2), indicating a higher pressure in the oocyte than in neighboring NC until stage 7 and a lower pressure in the oocyte from stage 8 onwards. Altogether, our data demonstrate that NC mechanical properties are graded along the antero-posterior axis at stage 8 and 9 because of pressure decreasing from anterior to posterior, preceding a possible increase in tension at late stage 9.

### The gradient of cytoplasmic pressure controls the wave of StC flattening

These data led us to hypothesize that the gradient of NC pressure could be responsible for the wave of the StC flattening. To test this, we used the *dicephalic* (*dic*) and *kelch* (*kel*) mutants, which have both been described as specifically affecting germline development [50, 51]. *dic* mutation leads to reduced germline growth [32] and *kel* mutations are known to prevent normal cytoplasmic transfer between the NC and the oocyte due to the presence of small ring canals (RC) [27, 51]. In *kel* mutants, the inner diameter of the ring canals fails to expand, leading to reduced lumen [52]. Based on these phenotypic descriptions, we expected that *dic* mutation would lead to a decrease of pressure whereas *kel* mutations would yield an increase in pressure. To prove this, we first analyzed NC membrane curvatures and observed that only half of the membranes bulge towards the posterior in *dic* follicles, whereas they are all convex in *kel* follicles (Figure 3A - 3C and Figure S5A). Similar results have been obtained for *Pendulin* follicles (data not shown). Mutations in *Pendulin*, which encodes a member of the Importin-alpha protein family, display small ring canals [53]. These observations imply that there is no gradient of NC cytoplasmic pressure in *dic* and that the gradient is more pronounced in *kel* and *pendulin* than in WT follicles. In the presence of the ROCK inhibitor, membranes in *dic* follicles still present convex and concave curvatures whereas in *kel*, most of the membranes remain convex. In parallel, no significant difference in Ecad or Sqh expression level along the A/P axis has been found between these mutants compared to WT (Figure S5B - S5D), confirming that cytoplasmic pressure, and not cortical tension, is mainly responsible for NC shape in these two mutants (Figure S5A). Second, AFM measurements show that the pressure is much lower in *dic* than in WT in S10 follicles whereas it is higher than WT in *kel* follicles. In *dic* follicles, the pressure gradient is no longer present (Figure 3D, 3F). In *kel* follicles, a difference between anterior and more posteriorly localized NC is still detected, but the values are more variable, as compared to WT (Figure 3E, 3F). These two mutants thus provide valuable mechanical conditions to determine the role of pressure and of the pressure gradient in StC flattening.

**Figure 3:**
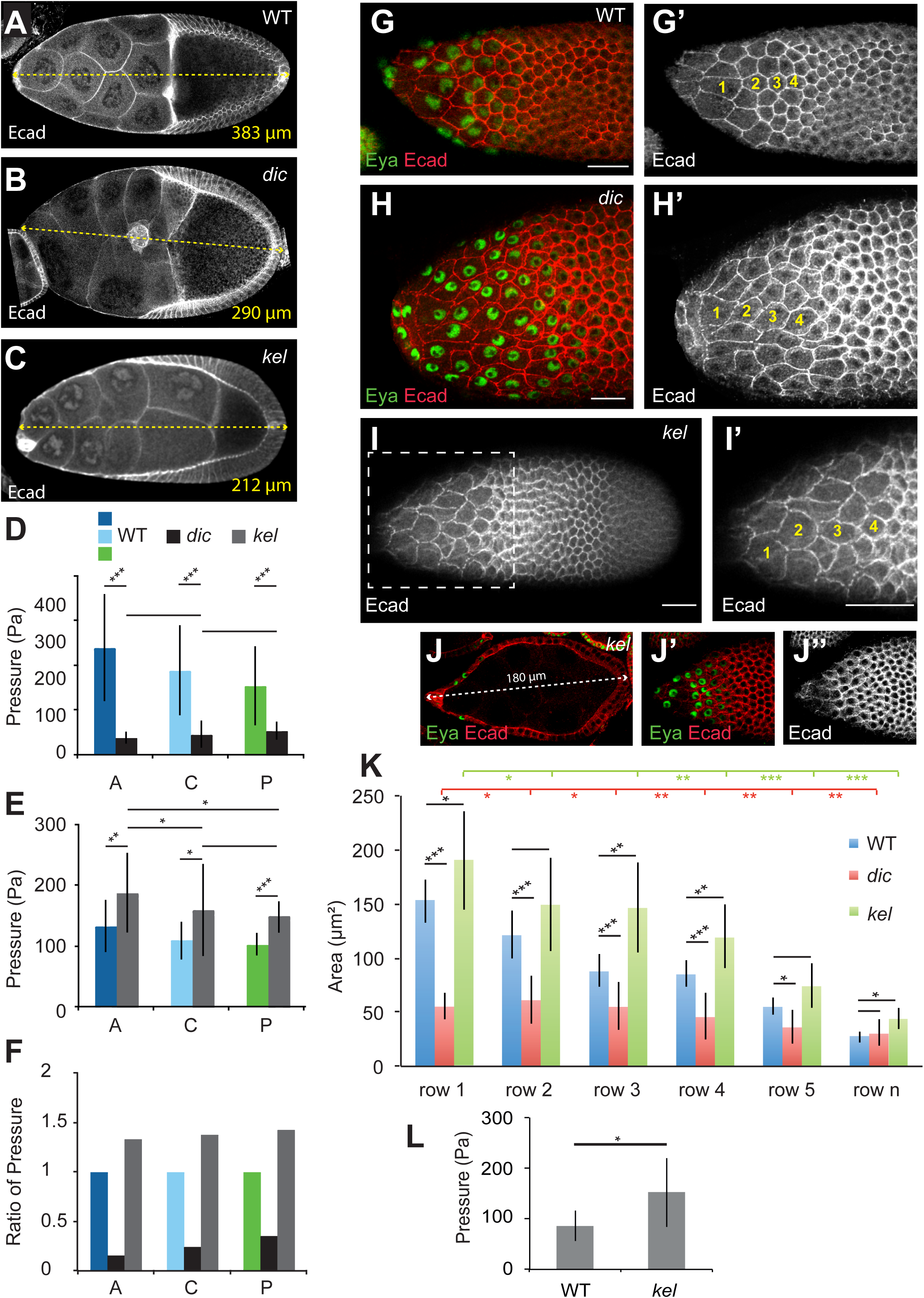
The gradient of NC pressure controls the timing and the wave of stretched cell flattening. (A-C) Cross-section of WT (A; stage 10), *dic* (B; late stage 9) or *kel* (C; mid stage 9) follicles showing nurse cell membrane curvature. (D) Inner pressure of S10 A, C or P NC between *dic* and WT (n > 15 for WT, n = 5 for *dic*). (E) Inner pressure of at S10 A, C or P NC between *kel* and WT (n > 10 for WT, n = 7 for *kel*). Note that WT values in D and E are specific to each experiment (see Methods). (F) Ratio of pressure in S10 A, C or P NC between *dic*, *kel* and WT. (G-H) StC flattening in mid S9 WT (G) or *dic* (H) follicles. (I, J) *kel* follicles displaying the progressive wave of flattening (I) or a premature StC flattening (J). Surface views (I, I’, J’, J’’) of the follicle or view across the follicle (J); I’ is a magnified view of the box drawn in I. For G-I, four cells are numbered in yellow along the A/P axis. For staging *dic* or *kel* follicles, see methods. (K) Apical StC surface area by position along the A/P axis in WT (blue), *dic* (red) or *kel* (green) mid S9 follicles (n > 10 cells for each row). Statistical comparisons between WT and *dic* or WT and *kel* are indicated in black, between *dic* rows in red and between *kel* rows in green. (L) Inner pressure of WT and *kel* anterior NC in mid S9 follicles (n=12 for WT, n=7 for *kel*). Scale bar: 20 µm.

The quantification of StC apical surface area shows that *dic* follicles start flattening later than WT and that no gradient of flattening is present (Figure 3G, 3H, 3K, and Figure S5E – S5H). In contrast, *kel* follicles present premature and gradual flattening, although it is less pronounced in the central area (Figure 3I - 3L, and Figure S5E - S5H). Together, these data demonstrate that the gradient of pressure controls the wave of StC flattening and that pressure levels dictate the timing of the flattening. It also confims that NC growth impacts the degree of StC flattening as previously mentioned [32].

### Inner pressure influences the temporal and spatial pattern of TGF**ß** signalling

We have previously shown that StC flattening depends on TGFß activity and that premature activation of the TGFß pathway is sufficient to induce early StC flattening [29]. The data presented above now demonstrate that StC flattening also depends on NC pressure and that contexts with increased pressure, such as in *kel* or *Pen* follicles, trigger flattening prematurely. To examine whether these findings are connected, such that the TGFß pathway in fact responds to NC pressure, we next monitored and quantified the levels of the phosphorylated form of Mad (pMad), a marker of TGFß activity, in WT and *kel* follicles. When follicle size is used as a benchmark to compare WT and mutant follicles, pMad is present as early as S6-7 in *kel* follicles (Figure 4A), whereas it is only detected from S8 onwards in WT follicles (Figure 4E). Additionally, pMad levels were usually two-fold higher in *kel* follicles than in the WT (Figure 4B - 4D). In both WT and *kel* follicles, the anterior and central areas display higher expression than the posterior area. These data show that high NC pressure is sufficient to induce TGFß signalling prematurely. In contrast, pMad is detected later in *dic* follicles, which display low NC pressure than in WT follicles (Figure 4E, 4G). Once expression commences in *dic* mutants, pMad expression levels attain similar levels to the WT, but is more uniform (Figure 4F, 4H, 4I). These observations show that pressure levels in the NC are important for the correct temporal and spatial pattern of TGFß activity.

**Figure 4:**
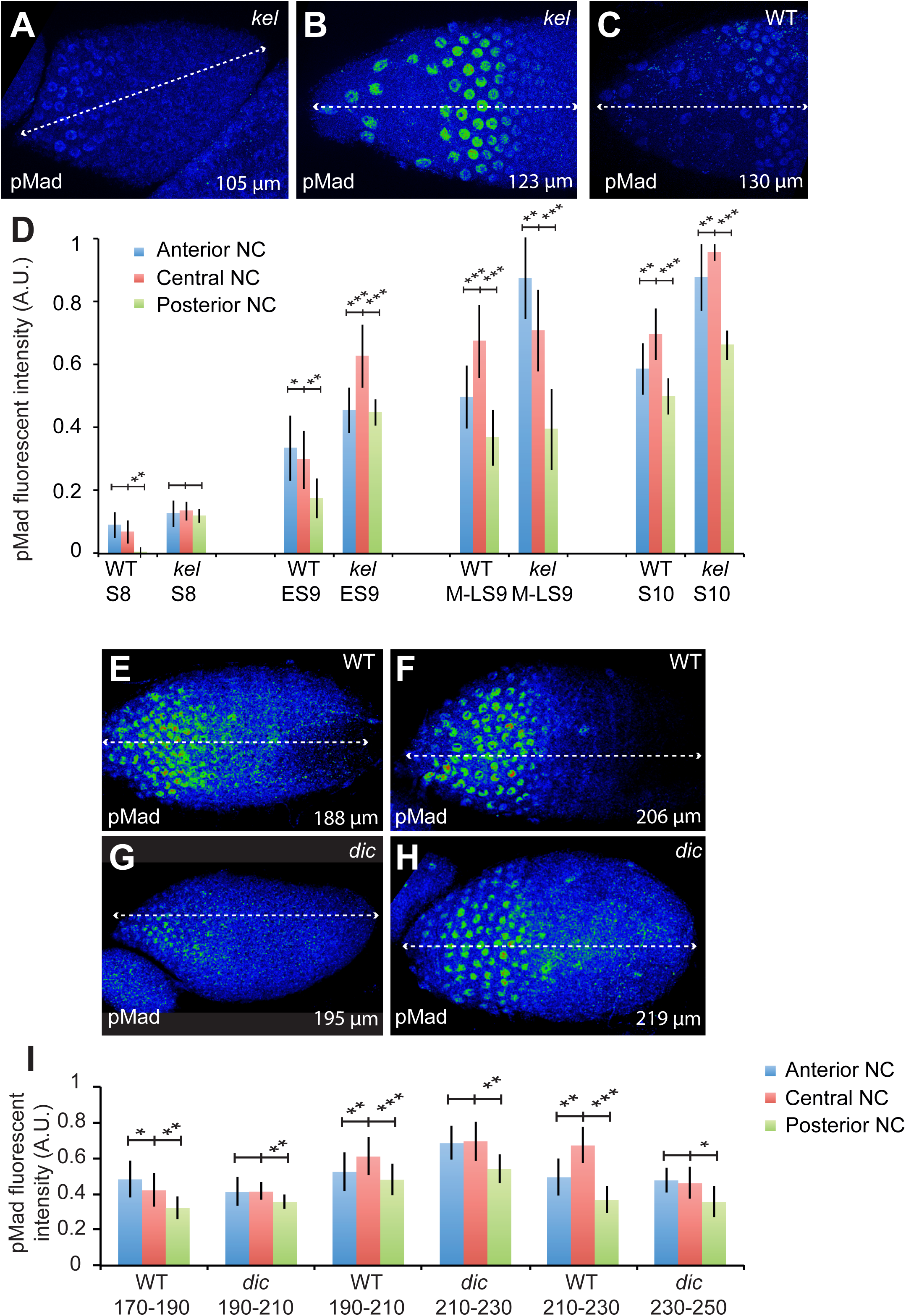
The pressure gradient controls the timing of the TGFß pathway. (A-C) Projections of the sections displaying pMad fluorescence in the StC in *kel* follicles (A-B) or in a WT follicle (C). (D) Quantification of pMad fluorescence in WT and *kel* follicles in function of follicle stage (n > 15 nuclei per region). (E-H) Sections through WT (E-F) or *dic* (G-H) follicles. (I) Quantification of pMad fluorescence in WT and *dic* follicles in function of follicle length (n > 15 nuclei per area).

### High cytoplasmic pressure, and subsequent StC flattening, leads to enhanced anterior follicle expansion

Our data show that anterior NC are more pressurized than posterior NC and the oocyte when follicles elongate. We reasoned that this might force the growing follicle to expand more anteriorly than posteriorly. To test this, we first used fluorescent beads that adhere to the BM and analyzed their movement during follicle growth (Figure S6A, S6B). From S8 to S10, we observed that beads at the anterior display a greater shift towards the anterior than posterior beads do towards the posterior (n=12) (Figure 5A, 5B and Figure S6C, S6D). Second, we used the *vkg::GFP* line, which expresses one chain of the Collagen IV fused to GFP, resulting in a fluorescent BM. We locally bleached the GFP contained in the BM in order to generate landmark points (n=5) (Figure 5C) and measured elongation of the anterior and posterior BM segments after 2 hours. On average, during WT S9 the anterior regions elongate 1.7-fold more than the posterior (Figure 5D), whereas at S7 or S8, no significant difference is detected (not shown). In stage 9 *dic* follicles, where pressure is decreased, the difference between anterior and posterior growth is reduced (n=6) (Figure 5D and Figure S6E). In parallel, we measured absolute follicle elongation and observed that *dic* follicles do not elongate as much as WT follicles, whereas *kel* follicles, where pressure is increased, are more elongated than WT (Figure 5E and Figure S6F – S6H). This suggests that NC pressure biases follicle growth towards the anterior during S9.

**Figure 5:**
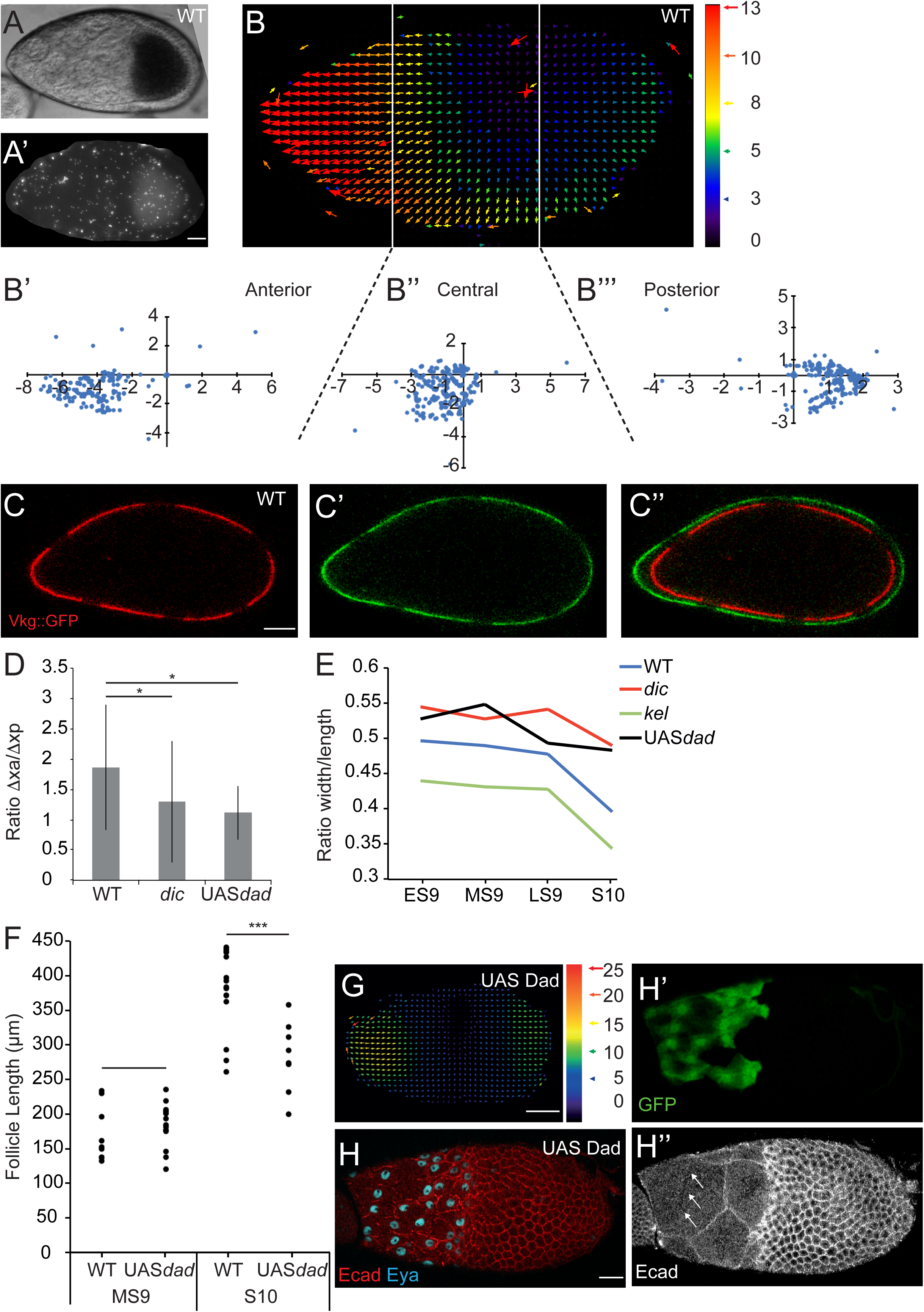
The follicles grow more anteriorly than posteriorly at S9. (A) A WT mid S9 follicle covered with fluorescent beads (A’). (B) Particle Image Velocimetry representation from the beads positioned on the follicle shown in A and plots presenting the coordinates of the vectors for the anterior (B’), central (B’’) and posterior (B’’’) areas. For each area, the initial (x, y) coordinates of the beads are (0,0). Each blue dot corresponds to the final x and y coordinates of a bead in µm. (C) WT mid S9 follicle from a female carrying the *vkg::GFP* transgene to mark the BM (red in C, green in C’). Six bleached areas are visible at t = o (C) and at t = 80 min (C’). The two images have been overlaid by aligning the central bleached areas (C’’) to show follicle growth during the interval. (D) Box and whisker plot of the ratio between the anterior (xa) and posterior (xp) growths of the BM in WT (n=9), in *dic* follicles (n = 6) and in Dad-expressing follicles (n=5). (E) Evolution of the ratio between the width and the length of the follicles from early S9 to S10 (n > 20 per stage). (F) Comparison of follicle length at mid S9 and at S10B between WT and Dad-expressing follicles (n > 20 per stage). (G) Particle Image Velocimetry representation from the beads positioned on a Dad-expressing follicle. (H) Mid-S9 Dad-expressing follicle. Only StC express Dad (GFP) (H’). The breakage of a NC membrane (arrows) are shown in H’’. Scale bar: 20 µm.

We then tested whether this anterior follicle expansion requires StC flattening and BM softening by analyzing follicle elongation when follicular cells constitutively express Dad, which represses TGFß activity and thus impairs StC flattening and BM softening [29, 31]. Our data show that Dad-expressing follicles indeed expand less anteriorly, are round and give rounder eggs than WT (Figure 5D - 5G and Figure S6I, S6J). These data show that anterior follicle and egg elongation requires TGFß activity in the StC.

One explanation for the lack of elongation in Dad-expressing follicles could be from the maintenance of a rigid BM in the anterior, mechanically preventing elongation. If this were true, one would expect that NC inner pressure to be high in Dad-expressing follicles. This is confirmed by AFM measurements that exhibit an increase of pressure by a factor of 1.4 between anterior NC from mid-stage 9 WT and Dad-expressing follicles (Figure S6K). Remarkably, we observed that Dad-expressing follicles display NC membrane breakage between the anterior NC at stage 10, suggesting that NC membranes collapse when pressure increases and anterior expansion is impaired (Figure 5H). This demonstrates that BM softening and StC flattening are important to maintain NC integrity, likely by promoting anterior expansion, which maintains NC pressure under a certain threshold.

### The establishment of the gradient of cytoplasmic pressure depends on intrinsic NC growth

As mentioned above, inner pressure varies as a function of differences in volume/growth rate. To precisely test the role of NC volume, we extracted 3D cell resolution data for each NC from S4 to S10 by using the MARS method (Figure 6A - 6F) [49]. Our data show that up to stage 8, volumes are rather similar between anterior and central NC, and that the oocyte is much smaller than any NC (Figure 6G and Figure S7A, S7B). From S8 onwards, the volumes of germline cells increase dramatically, especially of the four posterior NC and the oocyte (Figure 6G and Figure S7A, S7B). From S8 to S10, the ratio of the volumes of posterior and anterior NC is 2.4 ± 0.4 (mean ± SD) whereas the ratio of anterior and central NC volumes is only 1.1 ± 0.3. Our data show a gradient of volume between NC is present at these stages with the anterior NC being the smallest. No significant differences are observed between the central cells. The gradient flattens at S10, except for the four posterior NC. These data demonstrate that NC volume is regulated along the A/P axis during S9.

**Figure 6:**
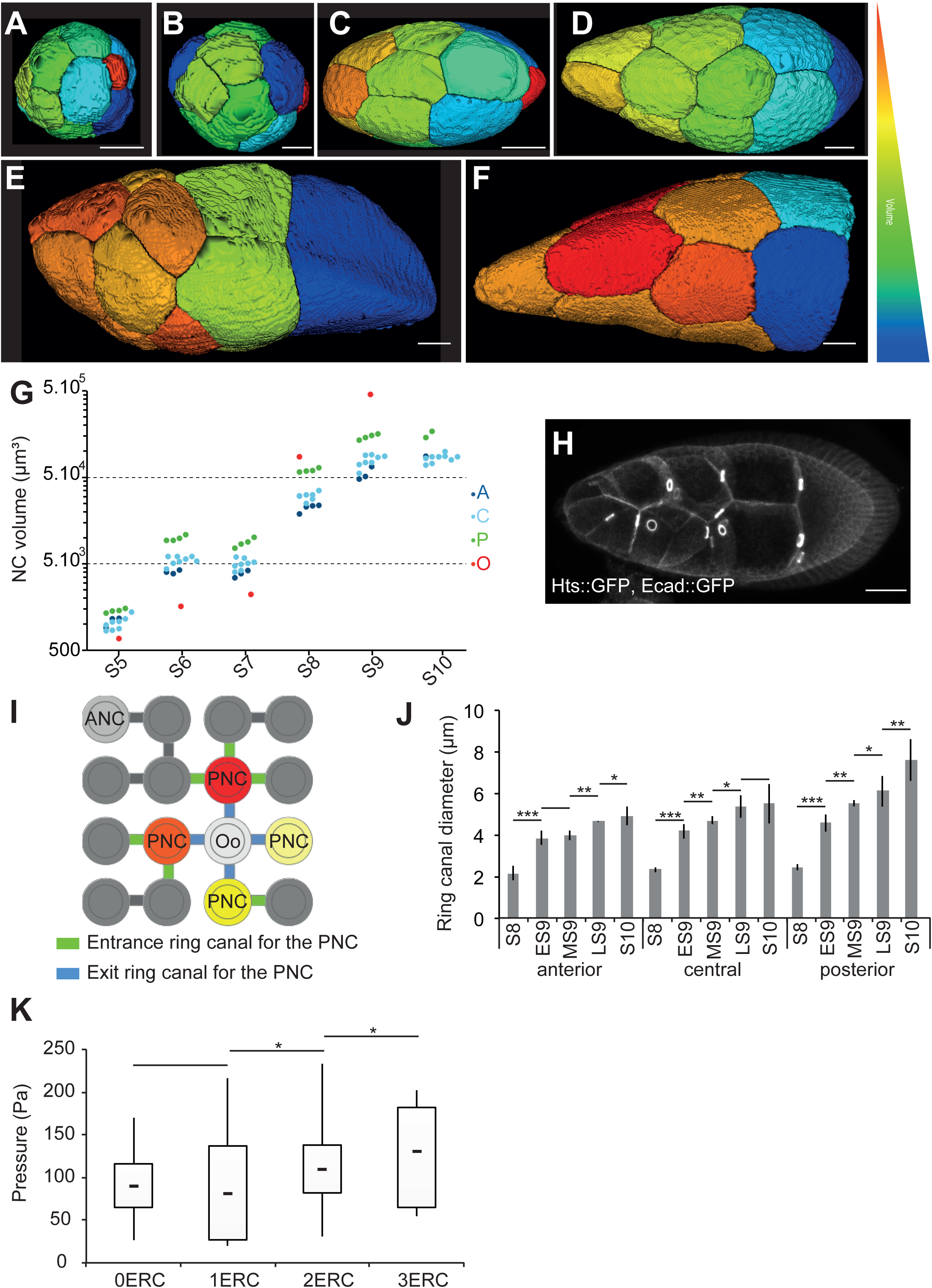
Intrinsic nurse cell growth is important for the pressure gradient. (A-F) 3D reconstructions of germline cells in WT follicles at S5 (A), S6 (B), S7 (C), early S9 (D), late S9 (E) and S10 (F). Only NC, but not the oocyte, are shown in F. The arbitrary color-scale indicates relative cell volumes for each follicle. (G) Individual volumes of anterior (A, dark blue), central (C, light blue) and posterior (P, green) NC and the oocyte (red) from S5 to S10 WT follicles. For S10, the segmentation was incomplete. (H) A S9 follicle showing the RC between the NC. (I) Schematic representation of the 16 germline cells and their stereotypic connections through RC The anteriormost NC (ANC) is shown in light grey. The four posterior NC (PNC, red or yellow) connect to the oocyte (Oo; white) via the RC shown in blue. One PNC (light yellow) has no ERC, while the three others have one, two or three ERC (green). (J) Quantification of RC inner diameters from S8 to S10 as a function of their anterior (A), central (C) and posterior (P) localisation (n = 5 per stage). (K) Box and whisker plot of inner pressure in the four posterior NC as a function of the number of ERC (n > 10 for NC with zero, one or two ERC and n = 5 for NC with three ERC). Scale bar: 20 µm.

NC volume depends on the invidual cytoplasmic content (the intrinsic growth), but also may depend on the transfer of cytoplasmic content from another NC through ring canals (RC) (Figure 6H). The four synchronous divisions that form the 16-germline cell cyst yield eight cells with a single RC, four cells with two RC each, and two cells with either three or four RC, with one of the latter being the oocyte (Figure 6I). To determine the importance of intrinsic NC contribution in building up pressure, we measured pressure in two NC with zero entrance ring canals (ERC) localized either in the anterior or in contact with the oocyte. The anterior NC always displays a pressure superior to the posterior NC (Figure S7D), indicating that intrinsic NC growth is an important parameter in controlling the NC volume, and therefore pressure.

### The gradient of cytoplasmic pressure is modulated by ring canals

According to Poiseuille’s law describing flow of a viscous fluid through a tube, cytoplasmic flux through an RC is proportional to the difference in pressure between neighboring NC and increases rapidly with the diameter of the RC. Our analyses of NC pressure in *kel* follicles, which have previously been shown to bear smaller RC [51], demonstrate that RC size plays a role in building NC pressure. Although it is known that RC undergo an approximately seven-fold increase in diameter throughout follicle development [27], it is unclear if any differences exist along the anterior-posterior axis. We have now carried out such measurements in the WT and confirmed that between S8 and S10, RC do indeed undergo progressive growth. In addition, we observed a gradient of size along the A/P axis, with the smallest RC in the anterior (Figure 6J and Figure S7C). Thus, RC size may participate in establishing the pressure gradient since narrower RC connect anterior NC, and wider RC connect posterior NC to the oocyte, which might help in building up pressure in the former and in preventing excessive build-up in the latter.

We then tested whether the number of entrance RC (ERC) per NC could also be related to different inner pressures. The oocyte has four ERC, each connected to a different posterior NC. Due to the pattern of cyst formation, the four posterior NC have either zero, one, two or three ERC [25]. Given what we have shown about cytoplasmic fluxes in the WT follicle, we hypothesized that the posterior NC with 3 ERC would have a higher cytoplasmic pressure than the one with no ERC. In 65% of cases, the highest pressure was measured in the cell that has the highest number of ERC (Figure 6K and Figure S7E), indicating that pressure at the same position along the A/P axis may vary as a function of the number of ERC. Importantly, the pressure in NC with three ERC is always lower than in more anteriorly-localized cells, which in most cases bear zero ERC. The number of ERC is thus not responsible for establishing the gradient, but likely only explains variations in gradient intensity.

## Discussion

Mechanical properties of biological elements are key components of organ morphogenesis, acting either directly by generating internal forces, or indirectly by applying forces and constraints on cells. Cell mechanical properties are usually considered to be determined by cortical tension. However, another potential contributor is cytoplasmic pressure, which serves to counteract cortical tension. Here we show that cytoplasmic pressure is graded within a well-defined group of nurse cells, and that this pressure then controls shape changes in the surrounding epithelial cells. These together sculpt the final organ, the egg, by indirectly acting on the mechanical properties of the basement membrane (Figure 7).

**Figure 7:**
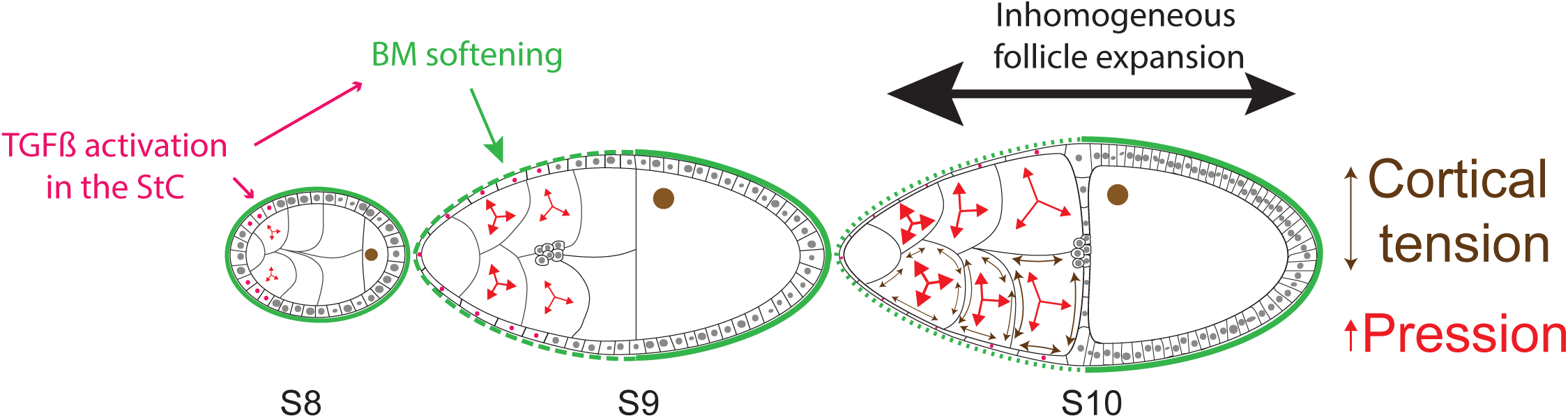
Model of the role of the NC cytoplasmic pressure on StC flattening and follicle elongation. From stage 8 to stage 10, a gradient of NC pressure (small to large red arrows) is established. When NC pressure reaches a certain level, it induces TGFß signalling in the surrounding StC (pink), allowing their flattening. The cortical tension (double sided brown arrow) appears to be inhomogeneous at late stage 9 and 10 (for simplification, it has been dawn at stage 10 in only half of the NC). In parallel, TGFß controls BM remodelling and softening (dotted to plain green line) [31]. This favours anterior follicle expansion (large black arrow) versus posterior expansion (small black arrow), allowing the growth of the NC. This growth allows the maintenance of the pressure, which continues to increase during stage 9 and 10, under a certain threshold, preventing NC membrane breakage.

The most direct method to measure pressure is through the use of a pressure probe, but a major drawback is that it does not allow multiple measurements within a single sample. Another possibility is to deduce pressure by measuring cytoplasmic flow between NC, either by observing the movement of cytoplasmic vesicles through ring canals or by photo-activating fluorescent molecules in one cell and observing their appearance in a neighboring cell. However, we were unable to efficiently detect vesicle movement through multiple ring canals along the A/P axis and photo-toxicity decreased the feasibility of performing multiple measurements within a single follicle. Therefore, we chose to measure pressure using a combination of non-invasive tools that do not compromise tissue integrity while simultaneously allowing us to test whether our measurements were dependent on cortical tension.

Our first approach, atomic force microscopy, makes it possible to probe deep into tissues in their native environment. Probing NC requires deforming basement membrane, StC, NC membranes and NC cortical actomyosin networks. Local cell integrity is preserved during experimentation, since no variation in pressure was observed over 100 measurements on a given cell. We show that our method is mostly sensitive to pressure by using different chemical treatments (NaCl treatment, collagenase) and genetic backgrounds (*dicephalic*, *kelch* and *Pendulin*). NC pressure values are in the range observed for other animal tissues, such as chick embryo blastula (10 to 200 Pa; Henkels et al., 2013), mouse lumen blastocyst (around 300 Pa; Dumortier et al., 2019), or for cultured cells (150 Pa; Stewart et al., 2011). In all three studies, pressure was similarly determined via non-invasive approaches, such as AFM or micropipette aspiration. Our second approach was to generate 3D reconstructions of the germline in order to infer differential inner pressure between the NC by measuring the curvature of their membranes [34], which allowed us to confirm the AFM measurements at late S9 and S10, and to establish that the gradient is already present at S8.

Sqh activity pattern suggests the existence of a postero-anterior gradient of cortical tension only at the end of stage 9, which is too late to influence StC flattening. Though gradients in both pressure and curvature persist in the absence of Myosin activity, treatment with the ROCK inhibitor results in a decrease in pressure values, indicating that actomyosin activity participates in NC mechanical properties. This could be due to the contribution of either the cortical or non-cortical network or both. It is not possible to evaluate the importance of cortical actomyosin activity between NC by laser ablation, since multiple ablations cannot be carried out in a single follicle. Overall, we conclude that NC mechanical properties are graded from stage 8 to stage 10 by cytoplasmic pressure with a possible late contribution of cortical tension.

While not all the components involved in establishing the pressure gradient are known, our data clearly show the implication of intrinsic NC growth, thus supporting the concept of proliferative pressure proposed as early as 1874 [54]. and of the size and pattern of the ring canals. By measuring precisely NC volume, we show that individual NC growth is not regulated as a function of pressure. It is known that DNA content of the NC is inhomogeneous at S9 and S10, with more DNA in posterior NC than anterior NC [26, 55]. Whether this difference in DNA content could participate in the establishment of the gradient remains to be tested. Our data also suggest that RC play dual and opposing functions in different NC: the large exit RC between the posterior NC and the oocyte may help prevent high pressure from building in the posterior, whereas the small RC in the anterior NC may help increase pressure there. The number of entrance RC also play a role in building pressure in certain posteriorly localized NC, as NC with such RC are mainly in the central and posterior areas [56].

StC flattening depends on both TGFß activity and germline growth (Brigaud et al., 2015, Kolahi et al., 2009). TGFß activity is required in the StC to set up the acto-myosin pattern that leads to AJ remodelling. *Mad* or *tkv* somatic clones lead to the presence of small and insufficiently flattened StC (Brigaud et al., 2015), indicating that NC pressure is not sufficient to induce cell shape changes in the absence of TGFß activity in the StC. By analyzing *dic* and *kel* follicles, we now demonstrate that NC inner pressure controls the triggering as well as the progression of StC flattening by acting on TGFß signaling. Nevertheless, a strict correlation between NC pressure and pMad levels during follicle growth cannot be drawn, possibly because once the TGFß pathway is activated by NC pressure, a negative feedback on the pathway, for instance through Dad activity, may kick in and lead to reduced expression of pMad, despite the continued presence of high NC pressure. It has been shown that Yorkie (Yki), a transcriptional co-activator, is present in the StC [57]. Yki is known to stimulate cell growth with DNA-binding partner proteins, including Mad [58], and to be regulated by cortical tension [59]. Future experiments will determine whether or not the molecular links between NC pressure and StC flattening requires TGFß and Yki.

Several studies have shown the importance of basement membrane structure and stiffness in the elongation of the follicle and the egg [60,61,70–73,62–69]. Two non-exclusive models have been proposed. First, that a softer basement membrane at both poles of the follicle would favor expansion along the A/P axis [74]. Second, that the structure of the basement membrane, with stiff fibril-like structures oriented perpendicular to the A/P axis, would prevent radial expansion [31]. The discovery of a pressure gradient within the NC reveals a third mechanism that acts from S8 to S10 (Figure 7). At S8, pressure increases in the anterior NC and induces TGFß in the StC, which in turn leads to flattening and to the local softening of the basement membrane. On the one hand, StC flattening likely facilitates NC growth, which is required to build up the pressure, by allowing a rapid transfer of oxygen and nutrients from the haemolymph. On the other hand, the softening of the BM above the StC likely helps the increase of NC volume and the maintenance of inner pressure under a certain threshold. Together, these intermingling genetic and mechanical regulations allow the coordination in growth and shape of two adjacent populations (StC and NC) that jointly act on the shape of yet another entity, the oocyte.

Our work highlights the role of fluid pressure in development, which has begun to be increasingly recognized [75]. Our results show that the ovarian follicle can be considered as a pressurized shell, with the basement membrane under pressure from NC and oocyte. This echoes experimental findings that adult *C. elegans* [76] and regenerating hydra [77] may be modelled as shells under pressure. We find that follicle growth is driven by cell pressure, similarly to regenerating hydra [77], and to elongation and inflation of the notochord in the Xenopus embryo [78]. It is worth noting, however, that cell-cell pressure differentials have been entirely overlooked so far. Our data show that an antero-posterior gradient in cytoplasmic pressure is important for follicle morphogenesis. One area that has been more explored is that of pressure caused by fluid accumulating within a lumen, such as in tubulogenesis during lung and kidney development [75]. Two recent studies reported the key role of lumen pressure in the mouse blastocyst in determining embryo size and cell fate, as well as in establishing the first axis of symmetry [48, 79]. More broadly, the implication of osmotic pressure in growth or morphogenesis is also well documented in walled cells such as in plants, fungi or bacteria [45, 80]. Importantly, some of these models also point out the importance of the balance between fluid pressure and extracellular constraints in sculpting organs, a balance that we also evidence in an animal system.

In summary, our work highlights the importance of the mechanical properties of cells or tissues neighboring a tissue undergoing a morphogenetic event and reveals the mechanical role of cytoplasmic pressure in shaping cells and organs.

## Methods

### Fly stocks and clones generations

Canton S was used as WT; the other fly stocks are: *hts::GFP* [81]*, Ecad::GFP* [82]*, PHPLC*γ*::GFP* [83]*, sqh::mCherry* [9]*, P(UAS-tkv^Q199D^)* (referred as TkvA)), *P(UAS-Dad.T*) [84], *traffic jam-Gal4* flies [85], *dic*^1^ [50], *kelch^ED1^* [51] *Pen^D14^* [53] and *vkg^G454^* (allele containing a GFP protein trap in the Col IV a2 chain Viking that we refer as Coll IV::GFP in the text, Morin et al., 2001).

Fly stocks were cultured at 25°C on standard food. Ectopic expression of TkvA or Dad were performed by generating Flip-out Gal4 clones in animals carrying the hs-FLP22 and the AyGAL4 UAS-GFP transgenes [87] or by crossing with *tj*-Gal4 flies. Flipase expression was induced by heat shocking 3 days-old females at 37.3C for 1 h to generate Flip-out clones. Adult females were fed on abundant yeast diet for 2 to 3 days prior to dissection.

### Follicle staining and staging

Ovaries from females were dissected directly into fixative 3 to 4 days after Flipase induction and stained following the protocol described in Grammont (2007). To avoid fluctuations of the depth of the follicles that are squeezed by the coverslip, each slide contains 15 ovaries, from which S11 to S14 are removed. After dissection of the follicles, most of the PBS is removed and 20µl of the imaging medium (PBS/Glycerol (25/75) (v/v)) is added before being covered by a 22/32 mm coverslip. The following antibodies were used: goat anti-GFP (1:1000; Abcam), rat anti-ECad (1:200; Developmental Studies Hybridoma Bank), rabbit anti-pMad (1:200; Nakao et al., 1997), mouse anti-Phospho-Sqh (1:1000; Cell Signaling).

The follicles were staged using their full length (Lf), the oocyte length for *dic* and Dad-expressing follicles, and the timing of the different morphogenetic processes occurring at S9 (border cell migration and stretched cell flattening) [25, 26].

### Fluorescent quantifications

Mutant and *viking::GFP*/CyO (which served as a control) flies were mixed before dissection. Both WT and mutant ovaries are thus fixed, stained and mounted together. Mutant and control follicles were discriminated thanks to the specific GFP accumulation in the basement membrane in *vkg::GFP* follicles.

For pMad quantification, a projection of all of the z sections in which StC nuclei are visible is made and background was subtracted. Nuclei were individually selected and registered as ROI. For each ROI, the ’’Area” and the “Integrated density’’ were quantified. Measurements were also performed in 3 areas located close to nuclei to estimate background and a mean of fluorescent background is calculated. The corrected total nuclei fluorescence (CTNF) for each nuclei is given by using the formula: CTNF = Integrated Density – (Area of selected nuclei X Mean of background fluorescence).

For Ecad, Sqh and pSqh quantifications, measurements were performed on the focal plan where the border cells are visible. ROI of 2 µm^2^ were positionned on plasma membrane based on the Ecad staining. The ROI measurement tool was then used to calculate the mean grey values for each area and the ratio was calculated.

### Atomic force microscopy (AFM)

AFM indentation experiments were carried out with a Catalyst Bioscope (Bruker Nano Surface, Santa Barbara, CA) that was mounted on an optical macroscope (MacroFluo, Leica) using an objective (10x objective, Mitutuyo). A Nanoscope V controller and Nanoscope software versions 8.15 and 9.2 were utilized. All quantitative measurements were performed using nitride cantilevers with silicon pyramidal tips (DNP-10 SCANASYST-FLUID+, Bruker AFM probes, Inc.) with a nominal spring constant of 0.7 N/m and a nominal tip radius of 40 nm. The actual spring constant of cantilevers was measured using the thermal tuning method [88, 89] and ranged from 0.6–0.9 N/m, which was sufficient to indent the sample without damaging it. The deflection sensitivity of cantilevers was calibrated against a clean silicon wafer. Fresh dissected follicles were fixed on a Petri dish coated with poly-L-lysine (0,5mg/ml) and were covered by living medium. Follicles are kept for an hour maximum before being discarded. All experiments were made at room temperature and the standard cantilever holder for operation in liquid was used. The Petri dish was positioned on an XY motorized stage and held by a magnetic clamp. Then, the AFM head was mounted on the stage and an approximated positioning with respect to the cantilever was done using the optical macroscope.

To record force curves, the Ramp module of the Contact mode in fluid was used. With this module, individual force curves are acquired at discrete points chosen using the optical image of the follicle. Each AFM measurement consists of the acquisition of 100 force curves extracted as 10 x 10 matrices with indentation points spaced 100 nm apart.

The pressure was deduced from local follicle stiffness and geometry [45, 90]. The local stiffness, *k*, is derived from the force–indentation curves by fitting to a linear model, using depths between 1 and 2 µm in order to probe the pressure of the NC while minimizing the influence of the basement membrane and of the stretched cells; the stiffness was obtained in piconewtons per meter (pN/m). The geometry at the indented location was characterized by the radii of curvature *R*_1_ and *R*_2_. Considering the follicle to be approximately a surface of revolution, *R*_1_ is the local radius of curvature of the follicle outline and *R*_2_ is the distance along the normal to the outline of the indented point to the axis of revolution. Radii were measured (in µm) from the optical images (from the macroscope) using ImageJ.

The pressure was then computed using the following equation [90]: *P = k*/π *2R*_1_*R*_2_/(*R*_1_+*R*_2_), with values in Pascals (Pa). The measurements referred to as “anterior” were not necessarily performed on the most anterior cell of the follicle, as the curvature of the follicle can sometimes prevent access. To reduce variability, each experimenter performed WT and mutant measurements and comparisons were made between sets obtained by the same experimenter.

### Bespoke protocol to probe internal NC pressure

Living follicles were dissected as previously described [31] and cultured in Poly-L-lysine coated petri dishes (Figure S1D). Measuring the apparent stiffness of NC requires deforming both the BM and the follicular cells (FC). We and others have previously established that the BM is on average 0.2 µm thick, whereas FC height varies from about 5 µm before S9 to less than 1 µm after flattening [31, 32]. Since AFM measurements typically allow a maximum of 3-4 µm indentation, we could not probe NC prior to S9 (Figure S1E). During S9 we probed one, two or three areas, located above the anterior, central or posterior regions of the NC, respectively, depending on the position of the wave of flattening (Figure S1F). No measurements were performed when StC height was superior to 1 µm. To be mostly sensitive to NC pressure, the stiffness of the cantilever and the force applied were chosen to enable indentation depths of 1.5 to 2.5 µm, which is more than the thickness of the BM and of the flattened StC combined. To perform the measurements, we used the Contact mode of the AFM, and registered 100 raw force curves from a 10 x 10 matrix, with indentation points spaced 100 nm apart.

### Chemical treatment of the follicles

Collagenase (1000 Units/ml CLSP; Worthington Biochemical Corp) was added to a final volume of 200 μl. The reactions were stopped with 10mM L-Cystein.

The ROCK inhibitor Y-27632 (Sigma-Aldrich) was added to a final concentration of 100 µM).

### Imaging for 3D-reconstruction

The MARS (Multi-angle image Acquisition, three-dimensional Reconstruction and cell Segmentation) pipeline is based on the fusion of several confocal stacks of images, taken with multiple angles, in order to recreate a highly resolutive three-dimensional reconstitution of the object. Segmentation is performed from the fused stack (for details, see Fernandez et al. (2010).

PHPLCγ::GFP follicles were dissected in PBS1X, before being stuck onto an coverslip coated with poly-L-lysine (0,5 mg/ml), which was glued on a Pasteur pipette. The pipette was fixed in a large Petri dish, allowing its rotation in three angles (-30°, 0 and +30°). Z-stacks were taken from the three angles using a 40X, 0.75 NA water-immersion objective of a Zeiss LSM700 confocal microscope (for S4 to mid-9 follicles) or using the 32X, 0.85 NA water-immersion objective and a pulse infrared laser (Chamaleon OPO) of a Zeiss LSM 710 (for thick follicles starting mid-late S9).

The follicles were either imaged in living conditions or fixed in a 4% formaldehyde solution and anti-GFP antibodies were used to detect PHPLCγ::GFP expression.

### Cell Curvature measurements

2D measurements are performed on fixed ovaries by manually applying circles fitting the curvatures between adjacent NC at a focal plane allowing border cell visualisation.

3D measurements are performed from segmented NC generated by the MARS method using a custom-made Image macro, called “Find-Curve”. The Find-Curve macro for the ImageJ program [91] automatically processes all cell stacks contained in a root folder indicated by the user. As the third dimension of the stacks, created by the MARS 3D reconstructions, corresponds to the Z axis, the program generates (X and Y) complementary orthogonal views. The three different views are then independently analysed. Using the “Default” threshold, derived from the Iso-Data algorithm [92], the macro identifies the slice with the largest area. An option has been implemented to manually specify the slice to analyse. On this slice, for every point (O) of the perimeter, the angle (OA, OB) is calculated, with A and B two neighbours (10^th^ degree) points, respectively upstream and downstream to O. The determination of local maximal angle values along the perimeter allows identification of the “summits” of the cell. These summits are used to define the various segments of the cell and generate the fitting regular polygon. For all segments, the radius of the best fitting osculating circle is calculated [93, 94]. This value is the curvature radius ρ of the studied segment. Finally, a .html report file is automatically created for human quality control.

A folder, called “test stacks for Figure 2G”, containing the three stacks corresponding to the nurse cells presented in Figure 2G and the macro can be downloaded by following this link: https://github.com/LBMC/FIND-CURVE

### Flow velocity measurements

Flies are dissected in PBS 1X and placed in a petri dish coated with poly-L-lysine and incubated 10 minutes with fluorescent beads (Fluoresbrite® Multifluorescent Microspheres 1.00µm 2,5% (Polysciences Inc.), dilution 1/100) before being rinsed and maintained in the living medium described in Montell. Images are acquired with Leica Macrofluo Plan Apo 5.0X/0.50 LWD objective. Stacks of images (z = 80 µm) were taken each 30 minutes and analysed with the ImageJ plugin, https://sites.google.com/site/qingzongtseng/piv [95]. The anterior, central and posterior areas are defined by dividing the length of the follicle.

### Basement membrane photobleaching

Homozygous flies for *vkg::GFP* were dissected and maintained in the living medium [96]. Follicles were placed in a coverslip coated with poly-L-lysine, which was glued to the bottom of a custom-made open chamber. Living medium was added from the top and covered by semipermeable membrane. Fluorescent images of the samples were acquired on an inverted Zeiss LSM 710 confocal microscope with 40x/0,75 water-immersion objective using the 488 nm line of an Argon Laser at 25°C. All images were acquired at a 512 × 512 pixels resolution. FRAP experiments were carried out by scanning over 20µm the follicle starting at about 15 µm from the coverslip. z-sections were performed with a step of 1 µm. The size of bleached region of interest (ROI) was about 10 x 10 x 10 µm, centred within the 20 µm-sized scanned area. Significant bleaching occurs after 20 iterations. All images were acquired at a scan speed of 4. Stacks of images were taken each 30 minutes over 2 hours.

### Statistical analyses

Normality of the samples was tested using Shapiro’s test. If at least one of the sample population did not have a normal distribution, the populations were compared with the non parametric Wilcoxon Rank Sum test. If both samples had normal distribution, their variance were compared using Bartlett’s test. If they were equal, a Student’s *t*-test was performed; if they were unequal, a Welch’s *t*-test was performed.

## Acknowledgments

We thank DSHB and Bloomington Stock Center for flies and reagents. We thank Lyon Multiscale Imaging Center’ (LyMiC) and Arthrotools of the SFR Bioscience (UMS3444/US8), as well as J. Chlasta for his help in adapting the MARS protocol to follicles and Steve de Bossoreille for suggestions. We are very grateful to A. Guichet, D. Bilder, Anne Ephrussi, J.R. Huynh, B. Loppin, Sally Horne-Badovinac for flies, V. Van de Bor for flies and discussion and F. Schweisguth for discussion. We acknowledge X. Wang for critical comments on the manuscript.

## Author contributions

Conceptualization (AB, MG); Methodology (PM, GR, PD); Software (AK, BV, PD, GR, DC); Validation (PM, PD, DC, AB, MG); Formal analysis (PM, DC, AB, MG); Investigation (LAL, PM, GR, LA); Writing – original draft preparation (LAL, MG); Writing – review & editing (AB, MG); Visualization (GR, PD, MG); Supervision (AB, MG); Project administration (AB, MG); Funding acquisition (AB, MG).

---

**Figure S1: Method for AFM measurements** (A-C) Sections through WT live follicles (A-B) or fixed follicles (C). (D) Diagram of the set-up used to perform AFM measurements. Living follicles are stuck to a Poly-L-Lysine coated Petri dish filled with culture medium. The AFM comprises a cantilever with a tip that is used to probe the follicles. The bending of the lever when it encounters the follicle is detected by a photodiode that captures changes in a laser-beam reflected onto the upper face of the cantilever. (E) Schematic representation of the probed area: the tip deforms the basement membrane (green) and the StC (red). In StC that have already flattened (B’), the underlying nurse cells (brown) are also deformed, allowing measurements. (F) Schematic representation of a late stage 9 follicle and of the cantilever: at this stage, only two regions were probed: above the anterior (A) and central (C) nurse cells. The posterior nurse cells are probed only in stage 10 follicles. For each region, a 10 x 10 matrix is measured, with indentation points spaced 100 nm apart. (G) Force-indentation depth curve obtained on a follicle with a pyramidal probe tip. The curve is fitted using the linear model (red line) to obtain the elastic modulus. Only the zone of interest (-1 to -2 µm) is fitted. (H) Schematic representation of the geometric measurements taken at areas where inner pressure was measured with AFM. Two circles are drawn to fit either the entire follicle (dotted black line) or of the AFM-probed area (dotted blue line at red dot), and the two radii, R1 and R2, are used to calculate inner pressure (see Methods). Scale bar: 20 µm.

**Figure S2: Control experiments for the AFM measurements** (A) Inner pressures of anterior (A), central (C) and posterior (P) nurse cells in individual WT follicles at stages 9 and 10. (B) Box and whisker plots of the 100 measurements taken in anterior (A), central (C) and posterior (P) nurse cells from a stage 10 follicle. (C) Color-coded representation of the 100 measurements in a single probed region of a WT stage 10 follicle. (D) WT stage 10 follicle before (D) and after (D’) NaCl treatment. (E) Force-indentation depth curves from A, C and P nurse cells of a WT stage 10 follicle before (solid lines) and after (dotted lines) NaCl treatment. Scale bar: 50 µm.

**Figure S3: methods for NC curvature in 2D and for NC 3D reconstruction** (A) Schematic representation of a WT stage 9 follicle representing the method used to measure nurse cell curvature in fixed follicles: circles (blue dotted line) are apposed to fit a particular nurse cell membrane and the radius (r) is measured. (B) Box and whisker plots of radii of the membrane curvature for A, C (both the anteriormost Ca and the posteriormost Cp) and P nurse cells in WT early stage 9 to stage 10 follicles (n = 98 cells for early stage 9; 42 for mid stage 9; 25 for late stage 9 and 15 for stage 10). (C) Percentage of convex (orange) and concave (blue) NC posterior membrane curvatures at different follicular stages in presence or not of the ROCK inhibitor (Y-27632). (D) WT follicle after addition of ROCK inhibitor. (E) Schematic representation of sample acquisitions for the MARS method. (F) 2D surface projections of a WT stage 8 follicle imaged at three different angles. Reference points (red dots) are used to fuse the stacks. (G) A mid-Z slice through a 3D reconstructed follicle where the three individual image stacks were fused into a single high-resolution stack. (H) 3D segmentation of the reconstructed follicle (G) showing only the germline cells. Scale bar: 20 µm.

**Figure S4: NC membrane curvature measurements from 3D reconstruction** (A) A mid-Z slice through a 3D reconstructed mid stage 9 follicle. (A’-A’’’) Segmented germline cells visualised from the z (A’), y (A’’) or x (A’’’) axis. (B) The stage 8 follicle presented in Figure 2G viewed at a different z section, along with the corresponding 3D segmentation (B’). B_A_, B_Ca_, B_Cp_ and B_P_ show to slices with the largest area in the 3D segmentation of the nurse cell under study, which are labelled as in Figure S2G. B’ panels present the outlines (solid white lines) of the same cells, as well as the circles (dotted white lines) fitting the posterior membranes (blue lines). (C) Percentage of anterior, posterior and lateral convex nurse cell membranes in WT stage 5 (S5) to stage 10 (S10) follicles. (D) Slice through a 3D reconstructed stage 6 follicle along with the 3D segmented image of its germline cells (D’). The slice with the largest area within the segmented oocyte in the z-axis is shown (D_Oo_) alongside its outline (solid white line) and the circles fitting the anterior membranes (blue line) (D’_Oo_ - D’’_Oo_). (E-G) Number of convex (orange), flat (grey) or concave (blue) curvatures of anterior nurse cell membranes (E), lateral nurse cell membranes (F) or anterior oocyte membranes (G) in WT stage 5 to stage 10 follicles. 2 or 3 follicles were reconstructed and segmented per stage. Scale bar: 20 µm.

**Figure S5: NC membrane curvature and StC flattening in dicephalic and kelch follicles** (A) Curvature orientation in WT, *dic* or *kel* follicle during stage 9 and 10 in presence or not of the ROCK inhibitor (Y-27632) (n > 10 follicles per stage). (B-D) Sections through *dic* live follicles (B-C) or *kel* live follicles (D). (E) Apical StC area along the antero-posterior axis, from row 1 (anterior) to row n (posterior), of WT (blue), *dic* (red) and *kel* (green) mid stage 9 follicles (n > 10 cells for each row). (F) Projection of all the sections where StC are visible in a stage 10 *dic* follicle. Most of the AJ are still visible (F’). (G, H) Apical StC area of row 1 (G) or row 2 (H), in function of WT (blue), *dic* (red) and *kel* (green) follicle length (n > 10 cells for each row). Scale bar: 20 µm.

**Figure S6: Particle image velocimetry representation for WT and mutants follicles.** (A) Late stage 9 follicle with fluorescent beads at two different time points (A, A1). A’ and A1’ are enlarged views of the yellow boxes drawn in A and A1, respectively. (B) The stage 9 follicle presenting in A. (C) Particle Image Velocimetry (PIV) representation from the beads positioned on the follicle presented in A. (D) PIV representation from the beads positioned on an early stage 9 follicle and plots presenting the coordinates of the vectors for the anterior (D’), central (D’’) and posterior (D’’’) regions. For each area, the initial (x, y) coordinates of the beads are (0,0). Each blue dot corresponds to the final x and y coordinates (in µm) of a bead. (E) PIV representation from the beads positioned on a stage 9 *dic* follicle. (F, G, H, I) Late stage 9 follicles from WT (F), *dic* (G), *kel* (H) or Dad-expressing (I) females with the length and the width of the follicles indicated. (J) Egg elongation from WT females or females expressing Dad in the follicular cells under the *actin* promoter (large Flip-out clones) (n > 40 per genotype). (K) Pressure in anterior nurse cells in WT and in Dad-expressing follicles at mid-stage 9. Scale bar: 20 µm.

**Figure S7: Global volume of the NC and difference of pressure between NC in function of the number of ring canals** (A) Volumes of anterior (A, dark blue), central (C, light blue) and posterior (P, green) NC and the oocyte (red) from S5 to S10 WT follicles. Central NC are further subdivided into those abutting the anterior NC (Ca) and those abutting the four posterior NC (Cp). (B) Volume of the oocyte from WT stage 5 to stage 9 follicles (n > 2 per stage). (C) Ratios of RC diameters at different follicular stages. Ratios are presented for anterior versus central (A/C) NC, anterior versus posterior (A/P) NC or central versus posterior (C/P) NC. (D) Ratio of pressure between the anterior-most (ANC) and the oocyte-connected posterior-most (PNC) single-ring canal (1RC) nurse cells (n = 4). (E) Inner pressure in two posterior nurse cells in WT stage 10 follicles. The number of entrance ring canals (ERC) of the two probed nurse cells is indicated by a symbol (diamond, triangle, circle or square).

**Movie 1:** Fused stack and 3D reconstruction of the germline of the stage 9 follicle presented in Figure S4A.

**Movie 2:** 3D reconstruction of the germline and the border cells of the stage 9 follicule presented in Figure S4A. The second part of the movie shows the border cells and the imprints of the posterior NC in the oocyte. The imprints are concave, indicating that the posterior nurse cells bulge toward the oocyte.

