## Supplementary figures and images for "Gradient in cytoplasmic pressure in the germline cells controls overlying epithelial cell morphogenesis"

### Fig S1

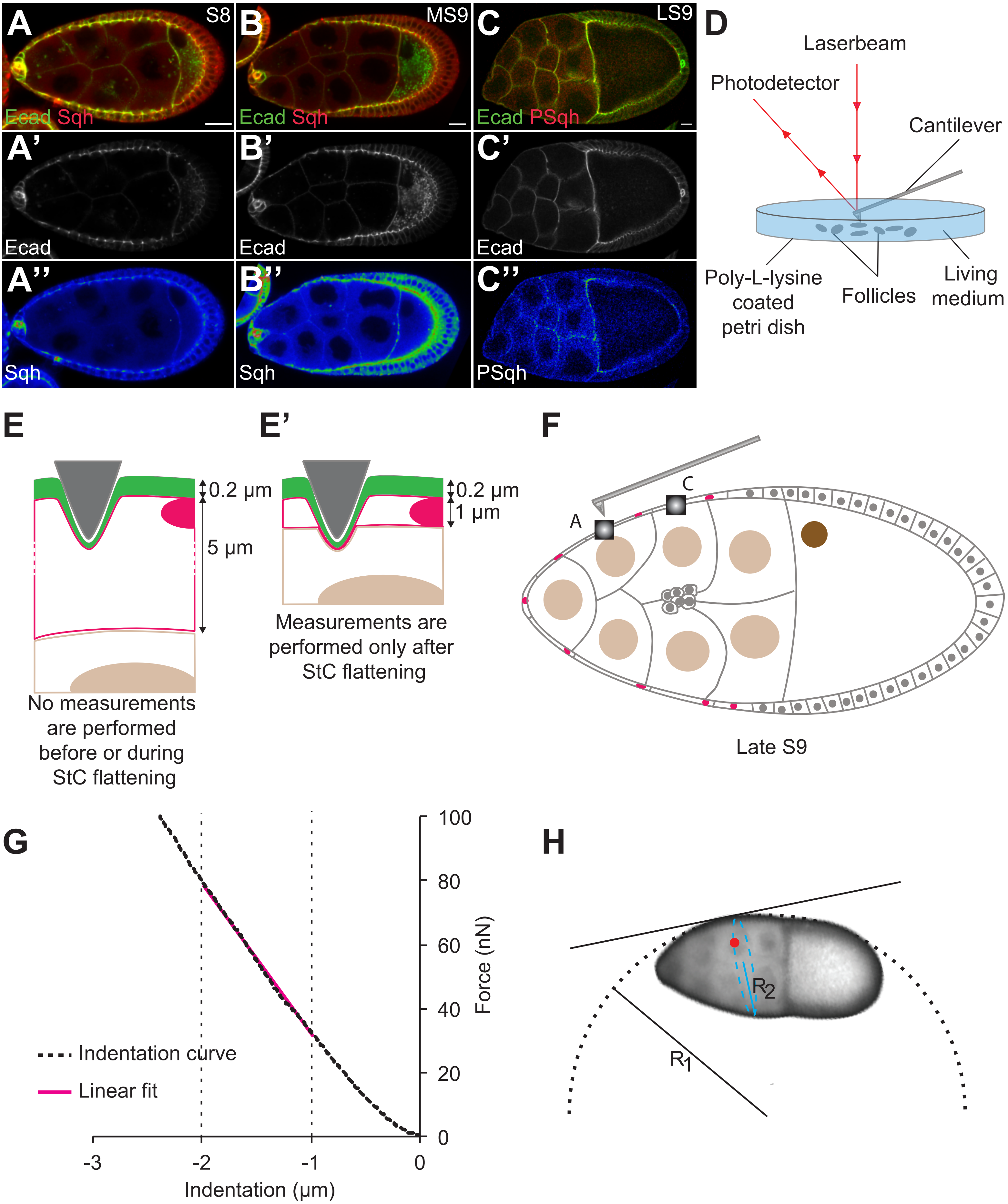

### Fig S2

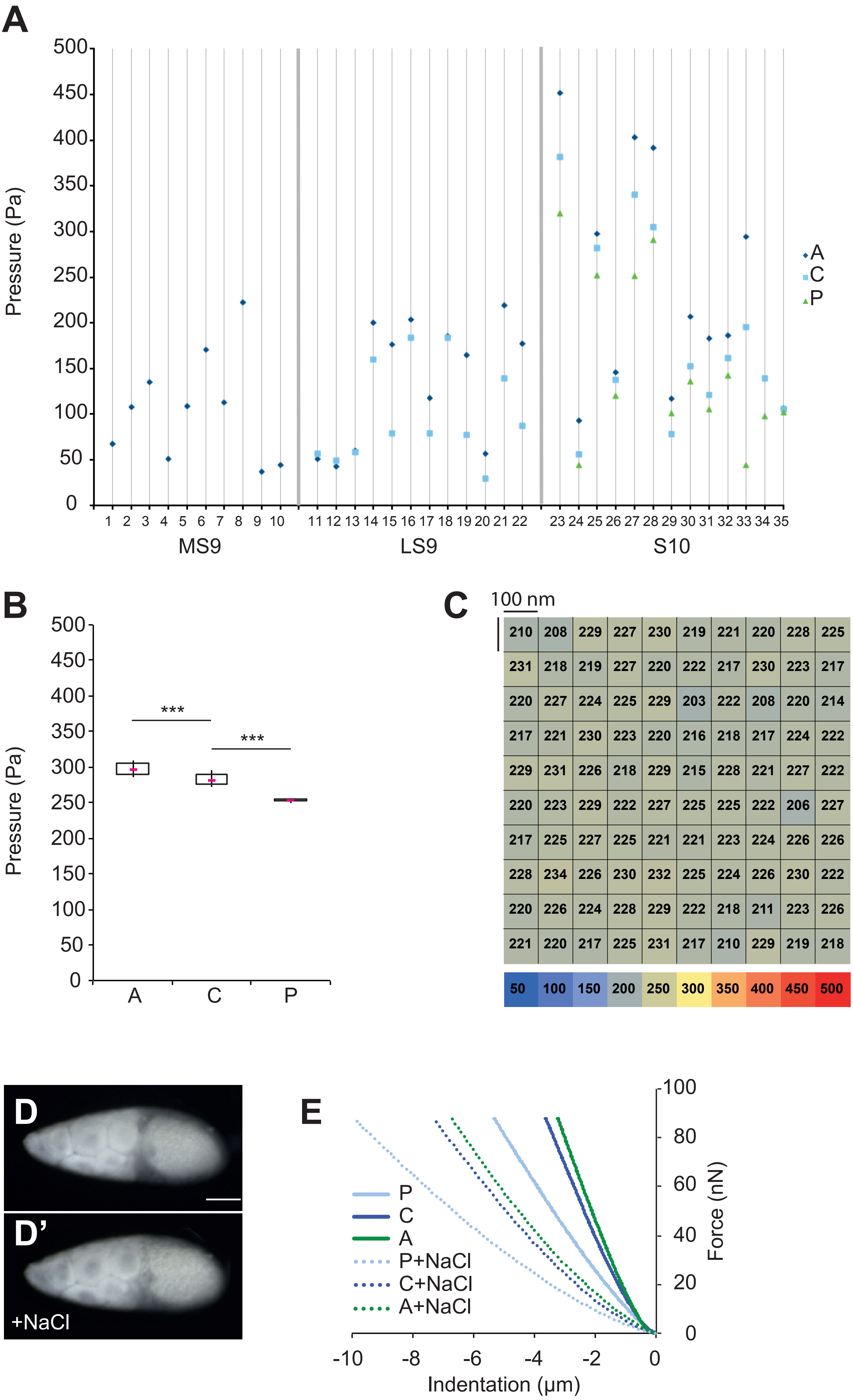

### Fig S3

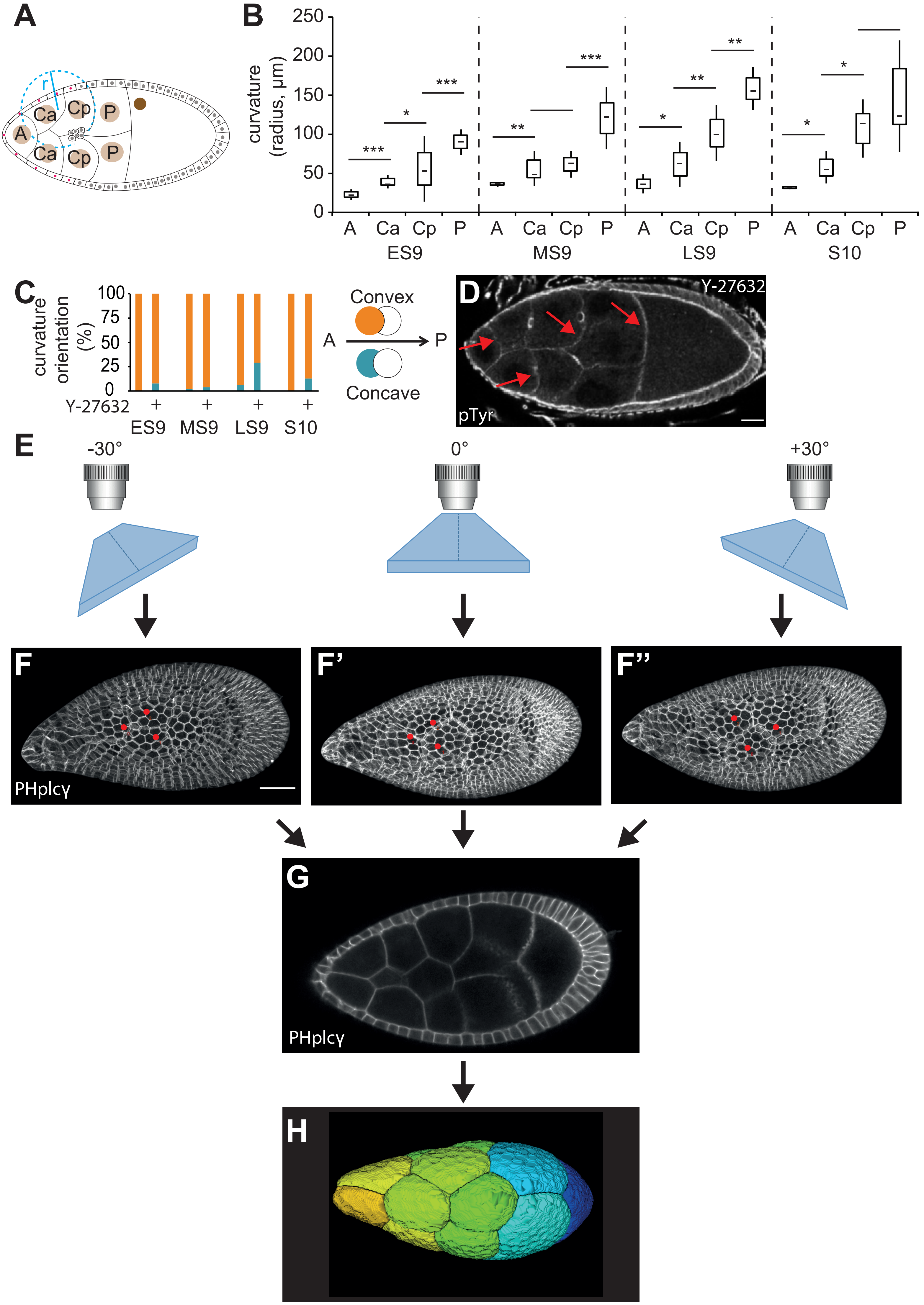

### Fig S4

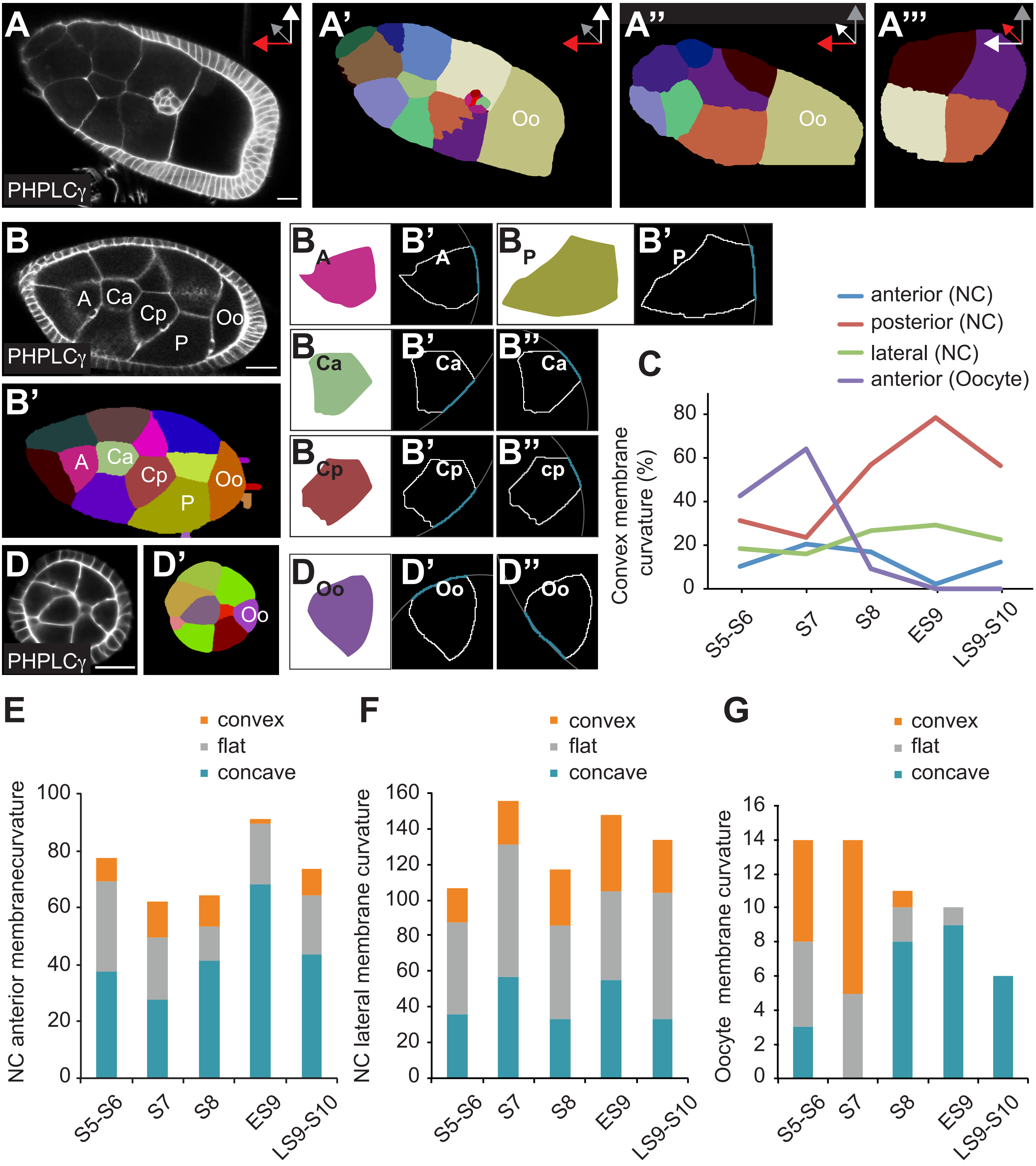

### Fig S5

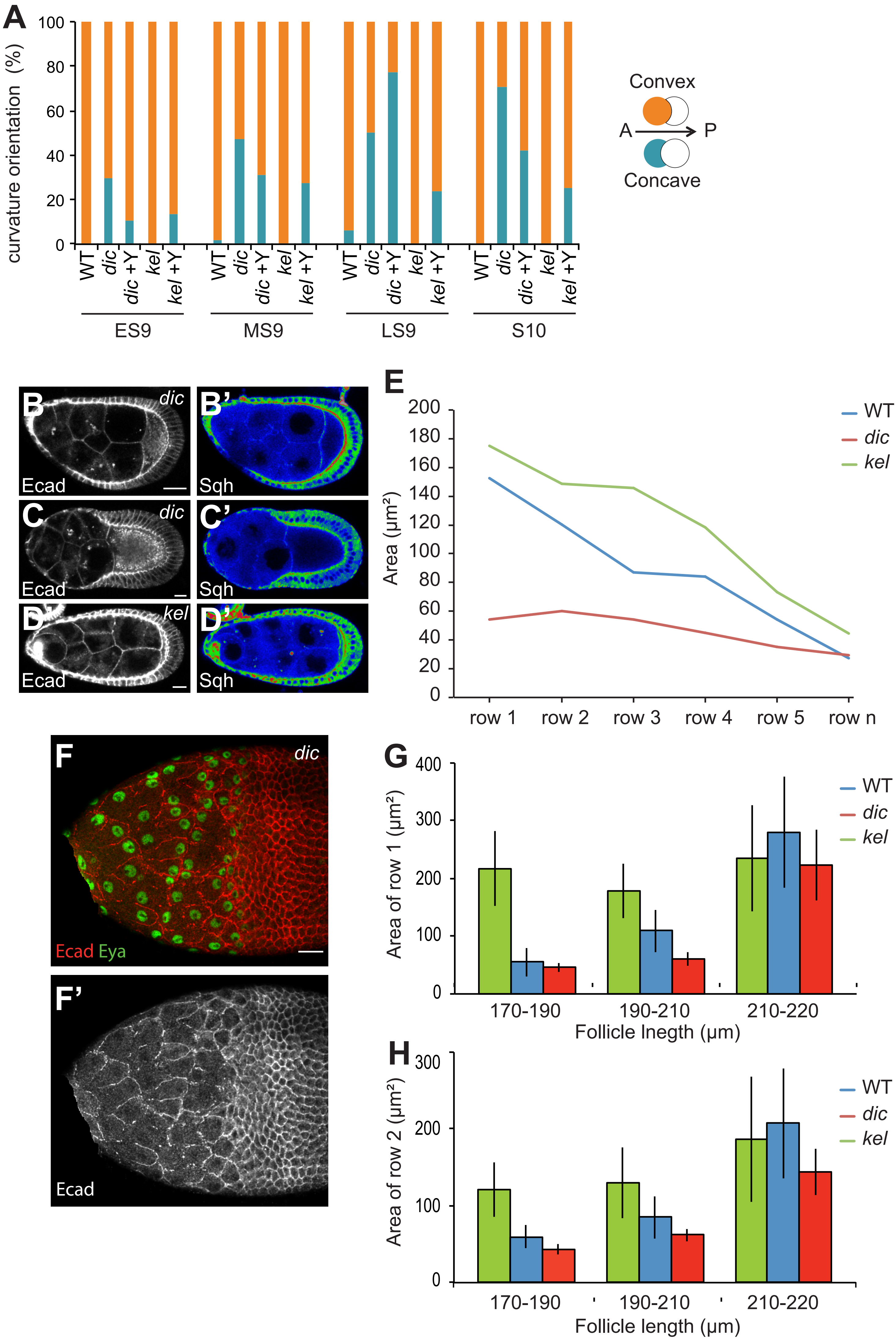

### Fig S6

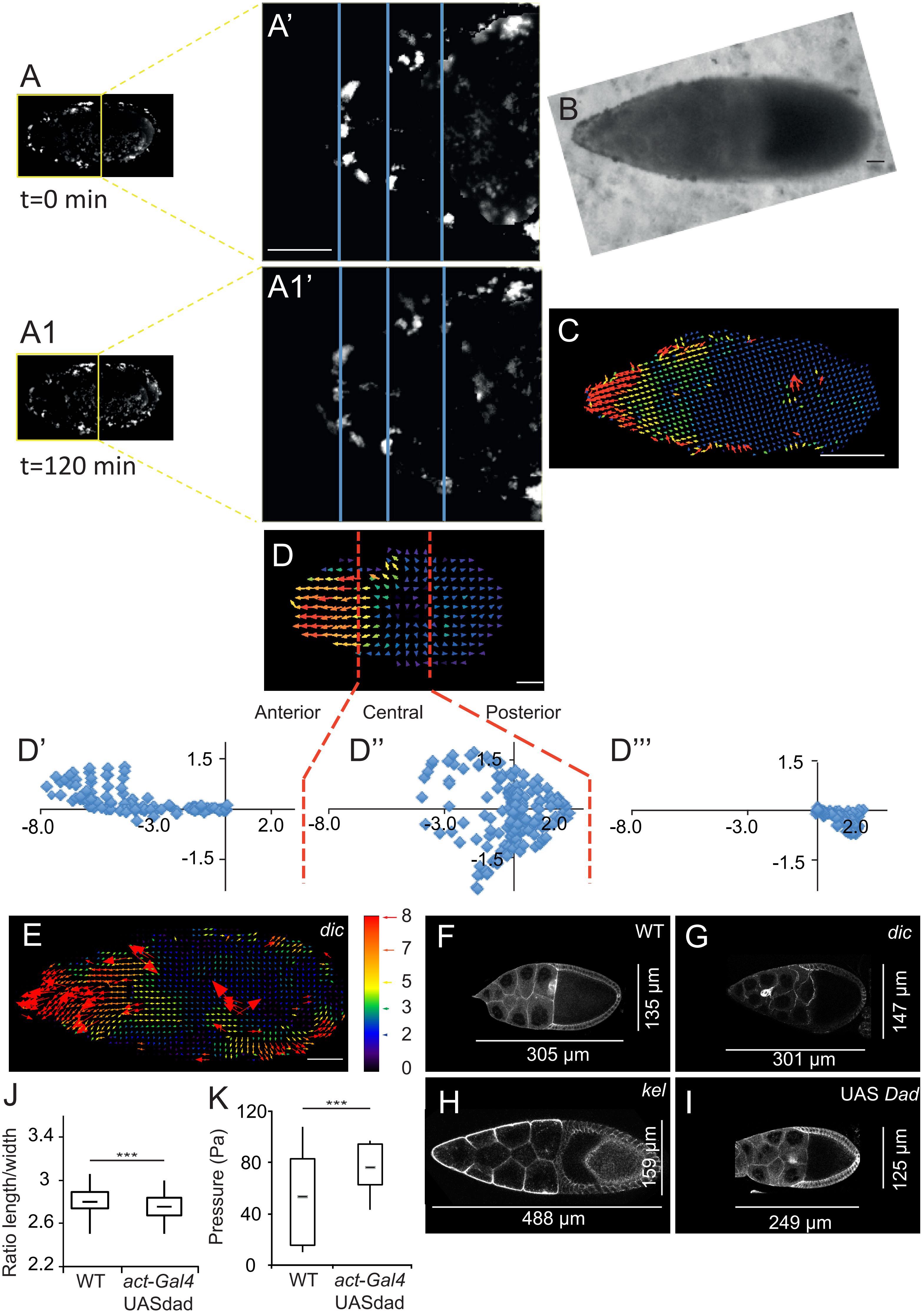

### FIg S7

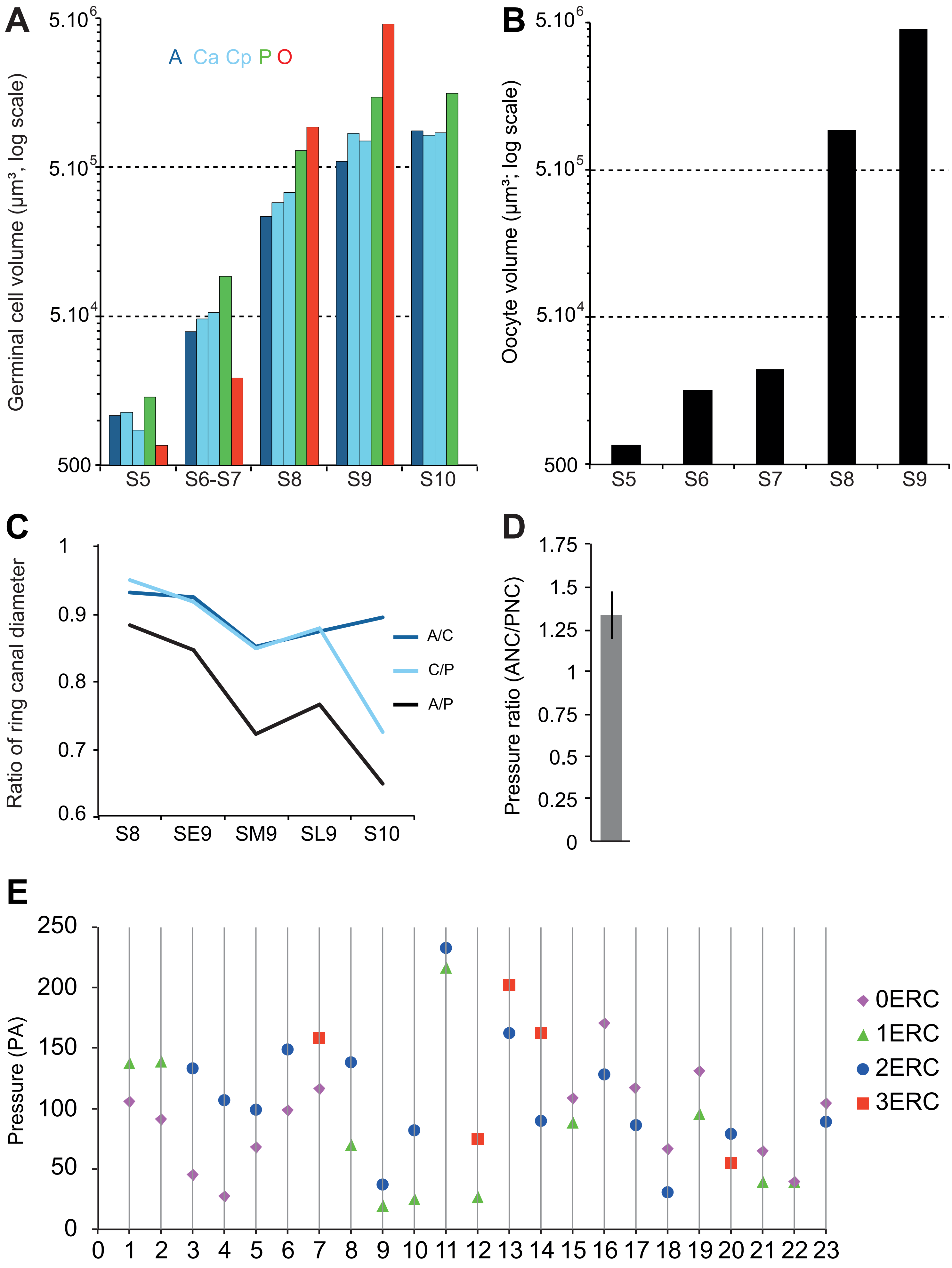
